# Self-organized tissue mechanics underlie embryonic regulation

**DOI:** 10.1101/2021.10.08.463661

**Authors:** Paolo Caldarelli, Alexander Chamolly, Olinda Alegria-Prévot, Jerome Gros, Francis Corson

## Abstract

Early amniote development is a highly regulative and self-organized process, capable to adapt to interference through cell-cell interactions, which are widely believed to be mediated by molecules. Analyzing intact and mechanically perturbed avian embryos, we show that the mechanical forces that drive embryogenesis self-organize in an analog of Turing’s molecular reaction-diffusion model, with contractility locally self-activating and the ensuing tension acting as a long-range inhibitor. This mechanical feedback governs the persistent pattern of tissue flows that shape the embryo and steers the concomitant emergence of embryonic territories by modulating gene expression, ensuring the formation of a single embryo under normal conditions, yet allowing the emergence of multiple, well-proportioned embryos upon perturbations. Thus, mechanical forces are a central signal in embryonic self-organization, feeding back onto gene expression to canalize both patterning and morphogenesis.

## Introduction

The regulative and self-organizing nature of the early amniote embryo is remarkably illustrated in experiments on avian embryos, in which the epiblast disk is subdivided into parts (*1*). Not only does the epiblast adapt to the absence of the removed parts, redirecting cell fates to eventually form a complete and well-proportioned embryo at its original location, but the other parts can spontaneously form additional, fully-formed embryos, albeit with variable frequency (*2*). The redirection of cell fates to form ectopic embryos has been shown to involve Gdf1 (also named cVg1), a TGF-β-superfamily secreted molecule that is normally restricted to the posterior side of the margin between the embryonic and extraembryonic territories; and whose ectopic expression at other locations along the margin accompanies the emergence of additional embryos in separated epiblast parts (*3*–*5*). Since Gdf1 is both necessary and sufficient to trigger embryo formation, a long-range, fast-diffusing inhibitor of Gdf1 emitted from the posterior has been postulated to explain the formation of a single embryo in intact epiblasts and the emergence of ectopic embryos upon subdivision, as the separated parts are freed from inhibition (*6, 7*). However, in a tissue that is millimeters across, it is unclear that interactions mediated by a diffusing inhibitor could support the rapid redirection of Gdf1 expression, downstream of the transcription factor Pitx2, itself detected as early as 3 hours following epiblast separation (*8*). And to date, a candidate for this role remains to be identified. Thus, although a number of molecular players have been identified, how the embryo self-organizes remains unknown.

We have recently shown that a supracellular actomyosin ring assembles at the embryo margin, and that its graded contraction (decaying from posterior to anterior) powers the large-scale rotational tissue motion that shapes the early embryo (Fig. 1A) (*9*–*11*). Thus, the entire margin is not only a molecular but also a mechanical organizer of development, where the shape and pattern of the embryo are simultaneously actualized. Here, using quantitative image analysis, mathematical modeling, and mechanical perturbation experiments, we investigate whether passive tissue tension, which propagates along the embryo margin in the intact epiblast (and is rapidly released upon cutting), could act as a fast-propagating and long-range inhibitor impinging on gene expression to control embryonic self-organization.

**Fig.1.**
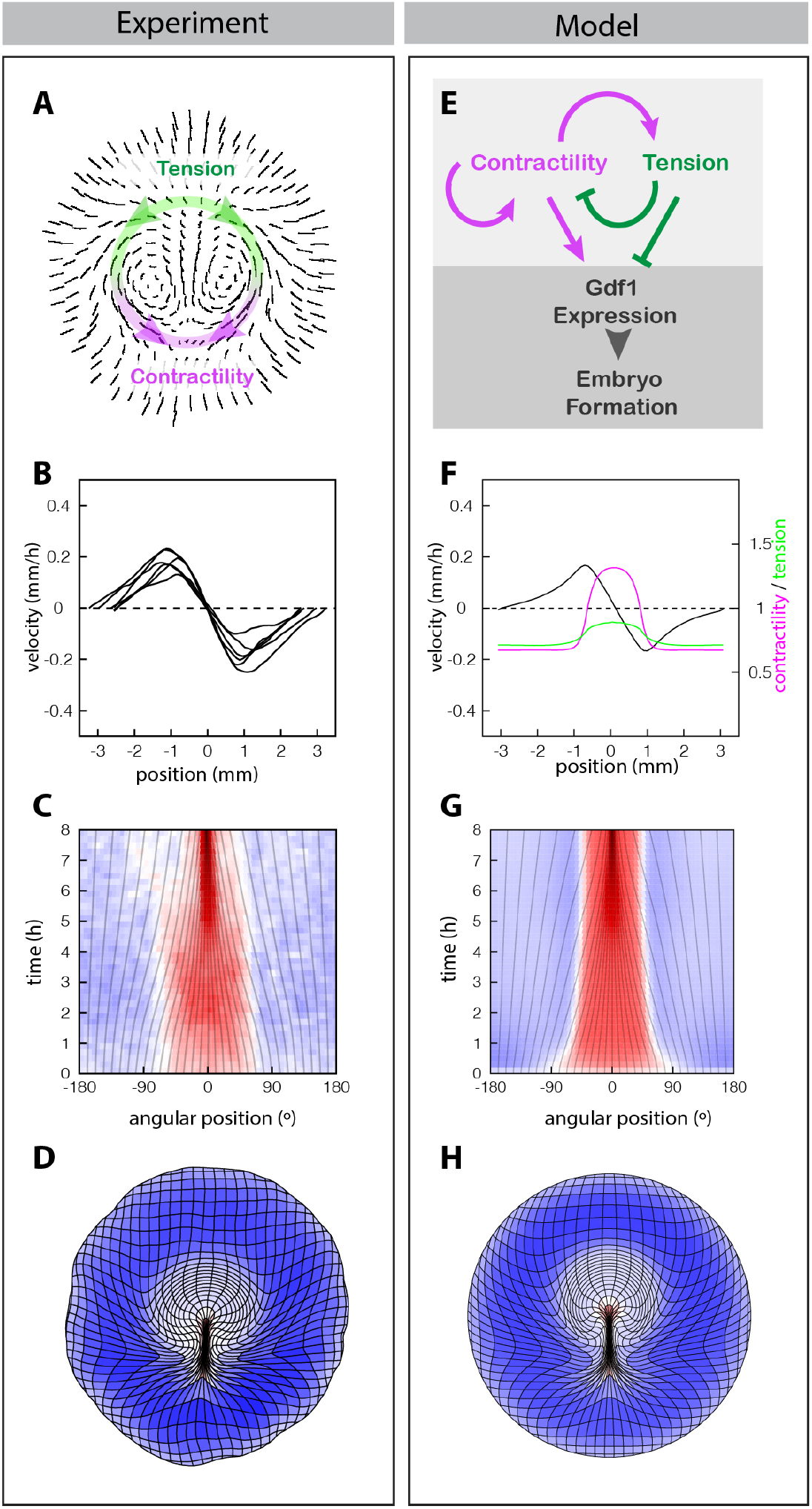
A mechanical analog of Turing’s reaction-diffusion model for the regulation of tissue contractility at the margin. (**A** to **D**) Trajectories [(A), t=6-8 hours; overlay denotes active contraction (magenta) in the posterior and passive tension (green) in the anterior margin], profile of velocity [(B), t=4h]) and time evolution of elongation rates along the margin [(C); grey lines denote the evolution of angular positions; 0° is posterior)], and deformation of an initially square grid (D), from an average of 6 embryos (11). (**E** to **H**) Interplay between active contractility (magenta) and passive tension (green) proposed to regulate embryo formation via Gdf1 expression (E) and predicted profiles of contractility (magenta), tension (green), and velocity (black) at t=4h (F) and time evolution of elongation rates along the margin (G), and global tissue deformation, when implemented in a in 2D fluid-mechanical model (H). Colors in (C, D) and (F, H) quantify contraction (red) and expansion (blue).

## Results

### A mechanical analog of Turing’s reaction-diffusion model for the regulation of tissue contractility

Our attention was drawn to the regulation of force generation in the embryo by the kinematics of tissue motion along the margin, as observed in quail embryos expressing a membrane-bound green fluorescent protein (memGFP) (*11*). Profiles of velocity and the time evolution of strain rates along the embryonic margin revealed a pattern of two domains of relatively uniform contraction in the posterior and stretching in the anterior (notice the triangular velocity profiles in Fig. 1B, with upward and downward slopes corresponding to stretching and contraction, respectively), which maintained stable proportions over time, even though the tissue continuously converges towards the posterior (Fig. 1C). The persistence of these domains over hours of development, even as critical regulators (Gdf1, but also Nodal, FGF8, and Wnt/PCP genes) are advected into the emergent streak (*5, 12*–*14*), hinted to a continuous regulation and to a degree of autonomy of mechanics from molecular signaling. To explore a possible role for mechanical feedback in embryonic self-organization, we formulated a minimal 1D model for the regulation of contractility at the margin, represented as a tensile line in which contractility locally self-activates and is inhibited by tension (Fig. 1E and methods). In effect, this is a mechanical analog of a Turing reaction-diffusion model (*15*), with contractility acting as a local activator and the tension it induces as a long-range inhibitor, and like a Turing model, it can support the spontaneous emergence and stable maintenance of domains of high and low contractility. In the embryo, tissue-scale contractility arises from alignments of cell-cell junctions into supracellular cables that are constantly turning over; the growth of these cables through the recruitment of new cell-cell junctions (as previously proposed (*16*)) may support local self-activation of contractility, whereas the passive tension that propagates all the way to the anterior margin could function as a long-range inhibitor by promoting their disassembly (as previously observed (*11*)).

To describe embryo formation, the same mechanical feedback was incorporated into a 2D fluid-mechanical model of tissue flows in the epiblast (*11*), with an initial bias in contractility representing the pre-existing polarity that directs embryo formation at the posterior margin (*5*). The model also allows for a nonlinear, saturating relation between the tension borne by contracting actomyosin cables and the rate at which they contract (cf. e.g. (*4*)), a refinement which we later show is required to account for the response to cutting (the model would otherwise predict a faster contraction in posterior halves as tension along the margin is released). The resulting model recapitulates the profiles of tissue velocity along the margin (Fig.1F, black curve), the maintenance of actively contracting and passively stretched domains with stable proportions, (Fig.1G), and the pattern of tissue motion entrained by the margin across the disk. Whereas our minimal 1D model is, mathematically at least, very similar to a molecular Turing model, this extended model identifies several effects specific to mechanics that play a critical role in the emergence of a single embryo. At odds with our minimal 1D model, in which tension propagates unopposed, in 2D its range is restricted by the ‘drag force’ from the surrounding tissue, and force transmission along the margin must dominate over propagation to the surrounding tissue to support inhibition in the anterior margin. Also, mechanical regulation must be fast enough to counteract advection towards the posterior and maintain a stable contractile domain.

With these effects accounted for (see supplementary text), self-organized contractility at the margin, implemented in a 2D fluid-mechanical model of gastrulation, recapitulates the full pattern of tissue motion that leads up to the formation of the primitive streak, the hallmark of the primary embryonic axis (compare Fig. 1D and H, see also Movie S1).

### Tissue contractility modulates Gdf1 expression to drive the formation of a single embryo in intact epiblasts

Our model, in which the balance between active contractility and passive tension along the margin governs the formation of a single embryo, predicts that modulating contractility locally, even transiently, should be sufficient to stably redirect tissue motion and form multiple embryos (Fig. 2A-F and Movie S2). Experimentally, these predictions were tested using beads soaked with Calyculin A or H1152, drugs that respectively increase and decrease myosin II activity. The model predicts that a transient inhibition of contractility in the posterior can split the posterior margin into two contraction foci (Fig. 2A-C); indeed, when H1152-soaked beads were placed at the posterior margin (Fig. 2G, H), we observed two sites of sustained contraction on either side of the midline (Fig. 2I). These contraction sites evolved into two primitive streaks, as revealed by tissue deformation (Fig. 2K) and Bra expression (Fig. 2J, L and Movie S2). Conversely, the model predicts that a transient increase in contractility in the anterior can give rise to ectopic contraction foci (Fig. 2D-F); indeed, Calyculin A-soaked beads deposited at the anterior margin (Fig. 2M, N) caused a local increase in tissue contractility and the emergence of ectopic contraction foci (Fig. 2O). These domains, as they contracted, were advected and often absorbed by the dominant contraction of the posterior margin, but occasionally elongated into additional Bra^+^ primitive streaks (Fig. 2Q, R and Movie S2).

**Fig.2.**
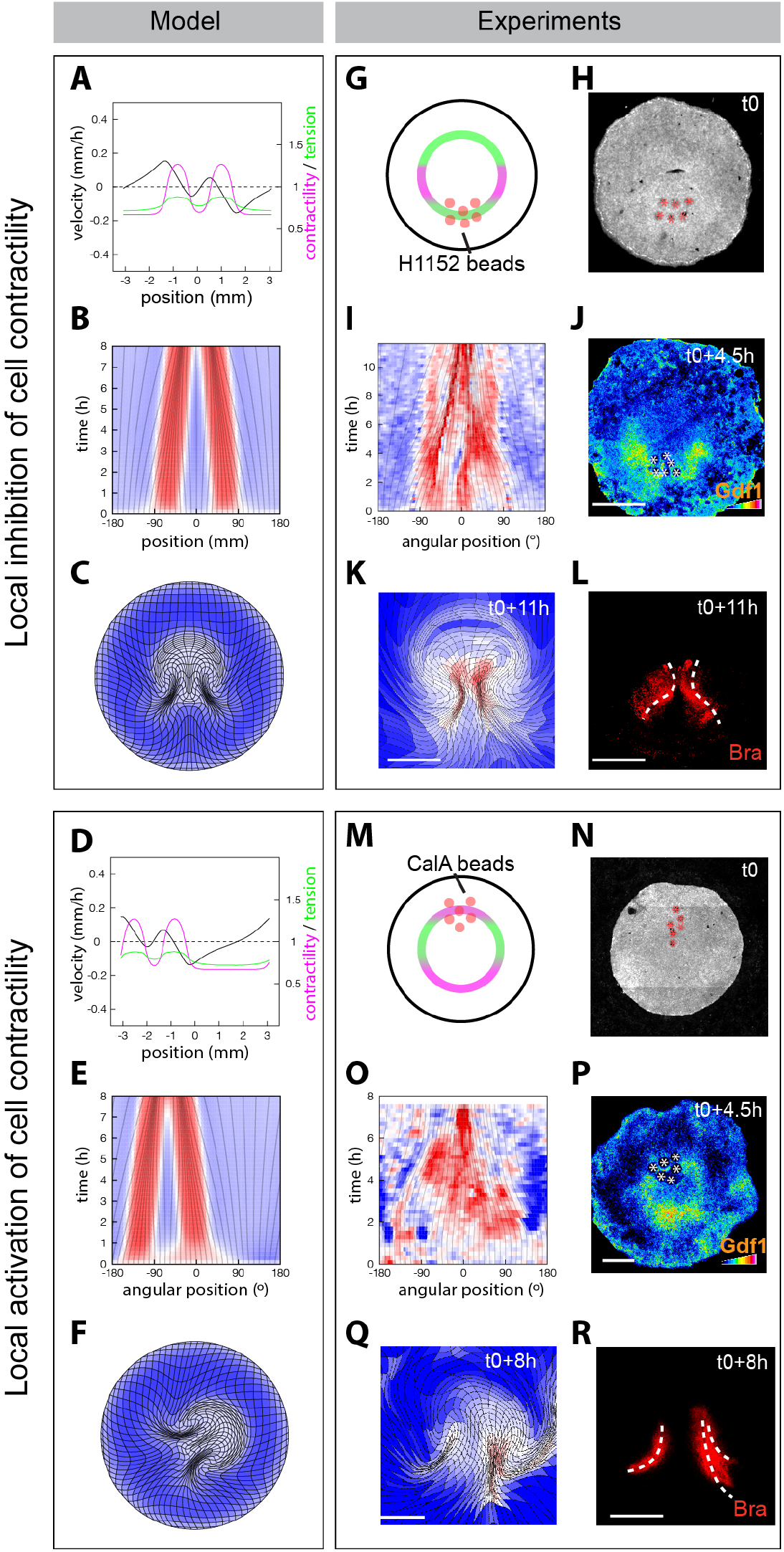
Tissue contractility modulates Gdf1 expression to drive the formation of a single embryo in intact epiblasts. **(A to F)** Model predictions for localized inhibition (A to C) and activation (D to F) of cell contractility at the margin. (A, D) Profiles of contractility (magenta), tension (green) and tangential velocity (black) at the margin; t=4h; 0 mm is posterior. (B, E) Time evolution of elongation rates along the margin; 0° is posterior. (C, F) Tissue deformation at t=8h. **(G** to **R)** Effect of locally perturbing tissue contractility by depositing of beads soaked in H1152 at the posterior margin (G to L) and Calyculin-A at the anterior margin (M to R). (G, M) Sketch of the experiment. (H, N) MemGFP embryo with beads (red asterisks) at t0. (I, O) Kymographs of elongation rates along the margin; 0° is posterior. (H, Q) Deformation maps and corresponding Bra expression (dotted white lines, primitive streaks; N=4/4 H1152 and N=3/5 Calyculin A). (J, P) Gdf1 4.5 hours after bead deposition (white asterisks, beads) (N=4/4 H1152 and N=4/5 Calyculin A). Scale bars, 1 mm.

Observing the expression of Gdf1 just 4.5 hours after the deposition of H1152 (resp. Calyculin A) beads, we found that it was locally downregulated (resp. upregulated) at the margin, consistent with the redirection of tissue motion (Fig. 2J, P). As controls, neither beads soaked in DMSO nor in H1152+Calyculin A induced ectopic Gdf1 expression or primitive streak formation in the anterior (Fig S1), demonstrating that these arise from specific effects on myosin II activity. Taken together, these results show that modulating tissue contractility at the margin is sufficient to redirect morphological and molecular hallmarks of axis formation via Gdf1 expression, and support mechanically self-organized regulation of Gdf1 expression as the key to the formation of a single embryo in the unperturbed epiblast (Fig. 1E).

### Self-organized tissue mechanics drives ectopic embryo formation upon epiblast subdivision

The classic epiblast subdivision experiments that revealed the regulative nature of early avian embryos have been so far interpreted in molecular terms (see the above-mentioned diffusing inhibitor model). However, cutting is a mechanical perturbation whose most immediate consequence is to interrupt the propagation of tension along the embryo margin (*11*). Our model predicts that in the absence of propagated tension, which normally acts as negative regulator, ectopic contraction foci spontaneously emerge. Assuming an initial posterior bias, as required to direct embryo formation in the posterior in the intact epiblast, two self-sustained contraction foci (and two streaks) are predicted to emerge at the posterior-most positions along the margin (Fig. 3A-E and Movie S3).

**Fig.3.**
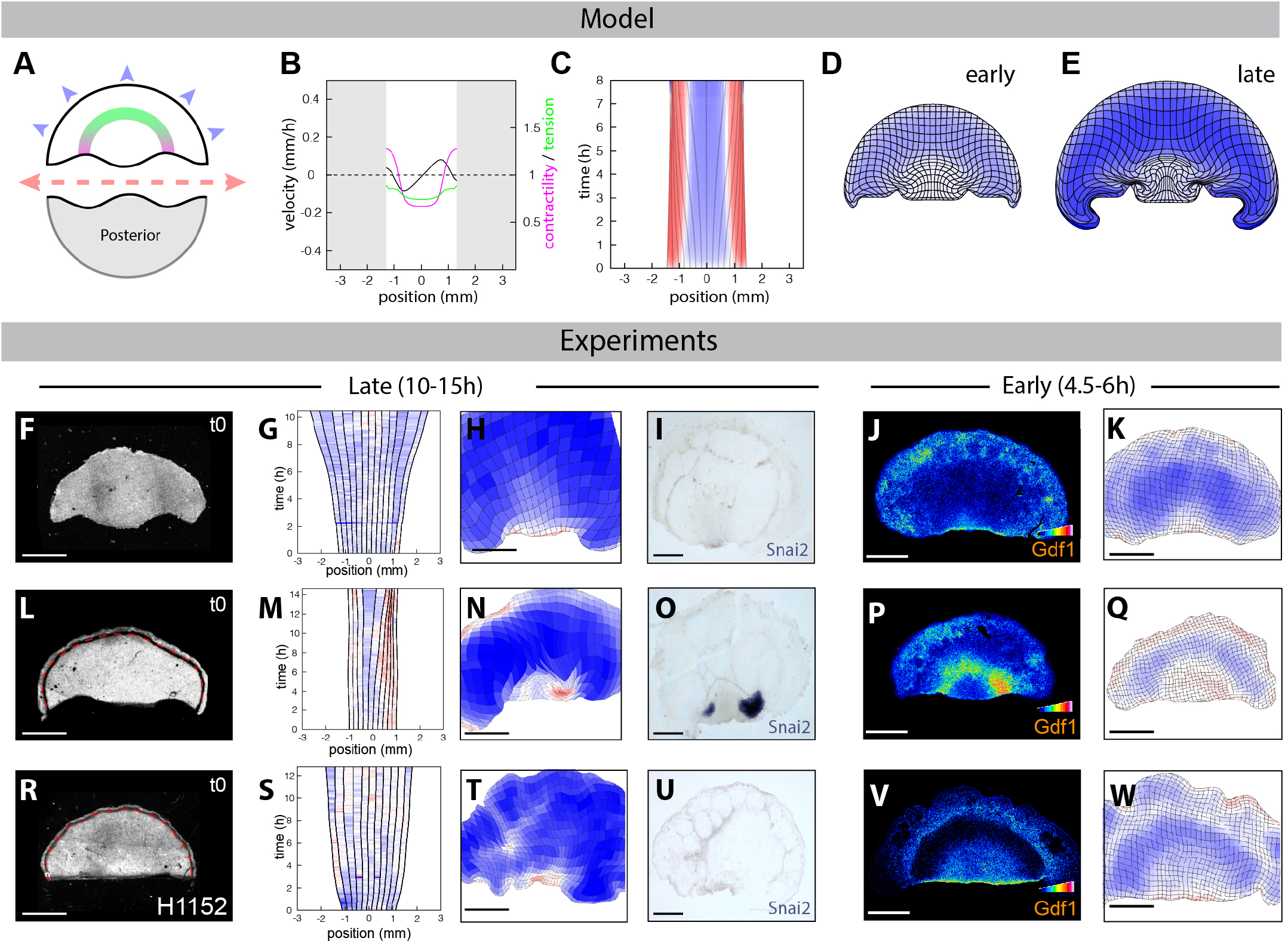
Self-organized tissue mechanics drives ectopic embryo formation upon epiblast subdivision. **(A to E)** Model predictions for the response to epiblast bisection in anterior halves. (A) Sketch of epiblast bisection. Predicted contractility (magenta), tension (green), and velocity (black) profiles at t0+4h (B), kymograph of margin elongation rate (C; 0° is anterior, and deformation maps at 4h (D) and 8h (E) after bisection. (**F** to **W**) UV-laser dissected anterior epiblast halves with (F to K) or without epiboly [(L-W); red dotted lines, UV cut abrogating the epiboly process)) and treated with H1152 (R to W). (F, L, R) MemGFP embryo at t0. (G, M, S) Kymographs of elongation rates along the margin; 0° is anterior. (H, N, T) Deformation maps at 10hours (H), 15 hours (N) and 13 hours (T) after epiblast bisection. (I, O, U) Snai2 expression in the same embryos fixed after live imaging [N=8/8 epiboly, N=5/5 without epiboly (3/5 show two ectopic primitive streaks, 2/5 one ectopic primitive streak), N=4/4 without epiboly+H1152]. (J, K, P, Q, V, W) Gdf1 expression and the corresponding deformation maps 4.5 hours (J, K and P, Q) and 6hours (V, W) after epiblast bisection. Scale bars, 1 mm.

To test a role for self-organized contractility in the redirection of Gdf1 expression and the formation of ectopic embryos in anterior halves, we revisited classic epiblast bisection experiments (*2*), with the added power of our quantitative analysis tools. Intact epiblasts were live-imaged for 1-2 hours, and tissue motion was analyzed in real time to locate the margin and presumptive axis (see Methods), allowing a precise bisection into anterior and posterior halves using UV-laser dissection. Anterior halves were then cultured for 4.5-6 hours (resp. 10-15h hours) to check for expression of early (resp. late) primitive streak markers (i.e. Gdf1 and Snai2; see Figure S2A-D and Movie S4). Unexpectedly, we did not observe ectopic primitive streaks in anterior halves in these conditions, based on either tissue deformation or Gdf1 and Snai2 expression (Fig. 3F-K and Movie S5). Ectopic primitive streaks only developed when epiblasts were bisected at an angle, such that the two halves had an overlap with the posterior domain of active contraction (Fig. S2 E-P). At odds with spontaneous symmetry breaking in the model, this suggested that the anterior margin may not support ectopic embryo formation in the absence of an activating trigger. However, analyzing tissue deformation in anterior halves, we noted that the margin and the embryonic territory as a whole were abnormally stretched (Fig. 3G, H, tissue expansion is in blue), compared to intact epiblasts or to biased anterior halves (Fig. S2). This suggested that the tension imposed on the embryonic disk by epiboly (the process by which the epiblast spreads via the migration of its edges on the vitelline membrane) might oppose the emergence of contractile domains at the embryo margin. Indeed, when epiboly was abrogated by uncoupling the migrating edge from the rest of the epiblast using laser dissection (Fig. 3L, red dotted lines and Movie S5), we observed the rapid emergence of contraction foci (within 1-2 hours) that expressed Gdf1 (by 4.5 hours) and eventually elongated into one or two Snai2+ primitive streaks (Fig. 3M-Q). Thus, the anterior margin does have the potential to initiate self-sustained contraction upon bisection, followed by Gdf1 expression and streak formation. However, this potential is susceptible to the counteracting effect of epiboly-induced tissue tension.

To corroborate a role for margin contractility as an upstream regulator of Gdf1 and primitive streak emergence upon bisection, we incubated bisected epiblasts with H1152 or Calyculin A, delivered uniformly in the culture medium. Whereas H1152 treatment prevented margin contraction, Gdf1 expression and subsequent primitive streak formation in epiboly-abrogated anterior halves (Fig. 3R-W), as well as in biased anterior halves (Fig. S2Q-T), Calyculin A treatment allowed margin contraction to overcome epiboly-induced stretching to form primitive streaks in anterior halves (Fig. S2U-X). Taken together, these results clarify the processes underlying classic subdivision experiments and demonstrate that upon physical cutting, redirection of tissue motion, through self-organized contractility, is the first event acting upstream of Gdf1 expression to drive primitive streak formation in anterior halves.

### Mechanical feedback (re)scales embryonic territories

While previous studies of embryonic regulation have given greatest attention to the emergence of ectopic embryos, our mechanical self-organization model also makes stringent predictions for the development of well-proportioned embryos from posterior halves. Indeed, the model predicts that the domains of active contraction and stretching adjust their proportions to tend towards a preferred, ‘homeostatic’ tension along the margin, shrinking if tension is too large or expanding when it is too low. It naturally follows that these domains should scale with margin length to restore homeostasis, as demonstrated in simulations of posterior halves (Fig. 4A-D). Critically however, this can only occur if the boundary of the tissue is fixed (if the cut edge of the epiblast reattaches to the vitelline membrane). If instead the boundary of the tissue is free, tension along the margin cannot build up again after it is released by cutting, and the actively contracting domain grows uninhibited to occupy the entire margin (Fig. 4I-L). By preparing posterior halves in different ways, we could favor or disfavor reattachment and challenge these predictions. As predicted, in epiblast halves that reattached, the contracting and stretching domains scaled down to occupy the same proportions of the half-length margin (Fig. 4E-G), so that the emerging primitive streak rescaled according to the new epiblast size (Fig. 4H and Movie S5). By contrast, in epiblast halves that failed to reattach, contraction spread to the entire margin and the primitive streak did not rescale (Fig. 4I-K). Of note, the margin did not contract faster in epiblast halves with free edges, implying that contraction is not limited by tension along the margin in the intact epiblast. (This justifies our choice of a model in which the contraction rate of cellular junctions tends to a plateau when contractility dominates over tension.)

**Fig.4.**
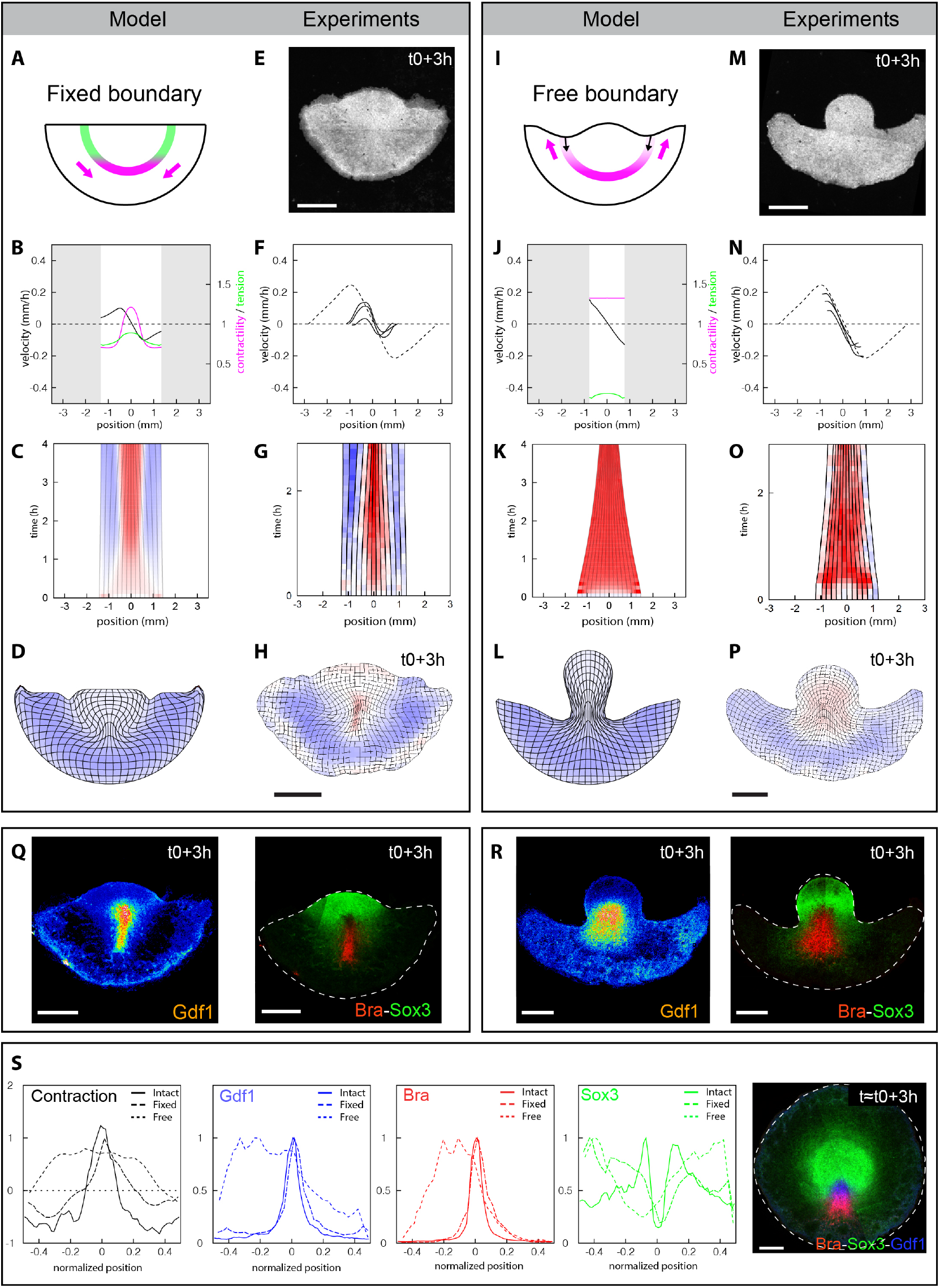
Mechanical feedback (re)scales embryonic territories. (**A** to **P**) Model predictions (A to D and I to L) and experimental response (E to H and M to P) to fixed and free boundary conditions in posterior epiblast halves. Sketch of the experiment (A, J) and memGFP posterior epiblast half after 3 hours (E, N). (B, F, J, N) Predicted and experimental profiles of contractility (magenta), tension (green) and velocity (black, dashed lines represent the averaged velocity profile of control intact embryos [N=5], shown in Fig. S3) along the margin at t0+2h. (C, G, K, O), kymograph of elongation rates along the margin. (D, H, L, P) Deformation maps at the end of experiments and simulations (t0+3-4h). (**Q, R**) Gdf1, Bra, and Sox3 mRNA in embryos shown in (E to H) and (M to P) respectively, fixed after live imaging. **(S)** Contraction and intensity profiles of mRNA signal along the margin in (Q and R) and compared to intact embryo (right panel; see also Fig. S3 for averages; the intensity profiles are normalized to the margin length). Colors in (C, D, G, H, K, L, O, P) quantify contraction (red) and expansion (blue). Scale bars, 1mm.

Having demonstrated a rapid redirection of tissue motion upon cutting that is controlled by mechanical boundary conditions, we asked whether this carries over to gene expression patterning. Indeed, just 3 hours after cutting, we observed a rescaling of the Sox3^+^ (neuroectodermal) and the Bra^+^/Gdf1^+^ (mesendodermal) territories (Fig. 4Q and S); quantification of Sox3^+^; Bra^+^ and Gdf1^+^ expression levels vs. relative position along the margin showed profiles that are strikingly similar to intact embryos (Fig. 3S). By contrast, in epiblast halves that failed to reattach, the Bra^+^/Gdf1^+^ mesendodermal territory expanded at the expense of the Sox3+ ectodermal territory (Fig. 4R, S). We note that in these experiments, stages of development were precisely quantified and matched between conditions to allow an accurate comparison (confounding effects such as might arise from the advection of gene expression territories can be ruled out; see Methods for details). Furthermore, when embryos with free edges were allowed to develop longer, the entire marginal tissue contributed to a fully elongated primitive streak, leaving a largely depleted embryonic territory to contribute to ectoderm, as opposed to posterior halves with attached borders, which maintain a balance between the pools of cells that contribute to different territories (Fig. S4 and Movie S6). Taken together, these experiments, in which only the mechanical boundary condition is changed, demonstrate that self-organized tissue contractility acts upstream of gene expression to enable the proper balancing and rapid rescaling of embryonic territories.

Mechanical self-organization provides a parsimonious account of the classic studies that initially revealed the regulative and self-organized nature of early avian development (*1, 2*). However, the physical cuts and separation they involve remain invasive perturbations, and one cannot rule out a contribution of wound healing or other responses elicited at cut edges to experimental outcomes. On the other hand, if these outcomes indeed manifest mechanical self-organization, it should be possible to achieve similar effects through less invasive mechanical perturbations interfering with tension propagation along the margin, such as localized friction. In order to introduce friction at the margin, a hair was deposited across the epiblast (Fig. 5A and 5E). The hair acted as an obstacle against the flow and slowed down tissue motion without provoking its complete arrest, indicating that it did not act as a tight barrier. As predicted by the model (Fig. 5B-D), two contraction foci rapidly emerged and elongated into primitive streaks in the anterior epiblast, just anterior to the hair (where tension propagation along the margin is obstructed), whereas the endogenous contracting domain in the posterior rescaled according to the new mechanical boundaries of the margin (Fig. 5F to H). The ectopic primitive streaks were preceded by the emergence of Gdf1 expression (Fig. 5I, K) and accompanied by redirection of Sox3^+^ and Bra^+^ embryonic territories by 4.5 hours after hair deposition (Fig. 5J, K).

**Fig.5.**
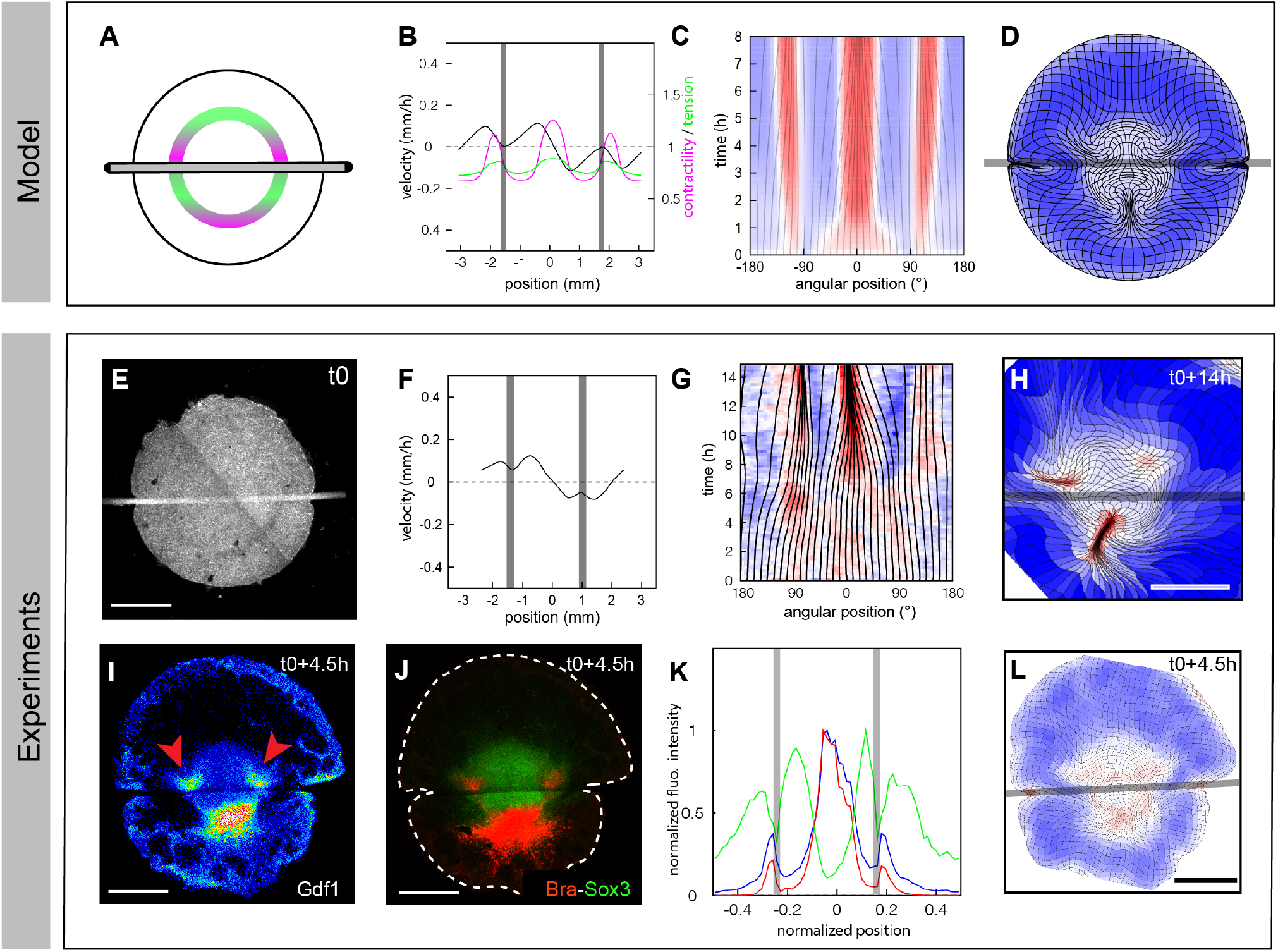
Mechanical friction redirects tissue motion/axis formation, inducing ectopic embryo formation and rescaling of embryonic territories. **(A to H)** Model predictions (A to D) and experimental response (E to L) to localized friction. (A) Sketch of the experiment and (E) memGFP embryo with a hair on its ventral side. (B, F) Profiles of contractility (magenta), tension (green) and velocity (black) along the margin at t0+4h; 0mm is posterior. (C, G) Kymograph of margin elongation rates; 0° is posterior. (D, H) Deformation maps at 8h (D) and 14h (H) after hair deposition. (**I** to **L**) Gdf1 (I) and Bra and Sox3 (J) expression, and corresponding intensity profiles along the margin (K) and deformation map (L), 4.5 hours after hair deposition (grey box). Colors in (C, G, H, L) quantify contraction (red) and expansion (blue). Scale bars, 1mm

Thus, mild non-invasive perturbations affecting the propagation of tension along the margin in intact epiblasts are sufficient to rapidly and persistently redirect force generation and tissue motion, leading to the emergence of ectopic embryos and the rescaling of embryonic territories via the modulation of critical gene expression.

## Discussion

In summary, our work uncovers the basis of embryonic regulation in early amniote development. While previous work focused on the molecular regulators of normal and ectopic embryo formation, we identify mechanical forces as a major signal in embryonic self-organization, upstream of gene expression. Because the actively contracting domain of the margin adjusts its size in response to these forces, embryonic regulation can be understood as a form of ‘mechanical homeostasis’. Homeostasis cannot be restored, and the embryo fails to rescale, if it cannot regain traction on the vitelline membrane following a cut.

The notion that mechanical forces could acts as a signal in embryonic self-organization has long been proposed (*17*–*19*). However, previous models remained theoretical and considered many possible effects, whose involvement during *in vivo* processes had remained difficult to disentangle. Furthermore, although it is becoming increasingly clear that self-organized developmental processes, such as gastrulation or feather follicle emergence (*20*–*22*), are sensitive to mechanics, a mechanistic framework accounting for the role of mechanical forces and tissue material properties, grounded in experimental observations, was still lacking. The interplay between active flows and patterning is perhaps most clearly worked out on the cellular scale, in early *C. elegans* development (*23*). In that context, advection of polarity markers that feedback on contractility is the key to establish embryonic polarity. A model that couples a bistable system with advection has also been proposed for avian gastrulation (*24*), but in that case, the time scale of advection is essentially the time scale of primitive streak formation, thus mechanical feedback that depends on advection cannot support rapid regulation, as revealed by subdivision (Fig. 4) and obstacle-based perturbation experiments (Fig. 5). By contrast, we propose that mechanical tension functions as a long-range signal that allows a near-instantaneous communication across the embryo to coordinate development. Provided that force transmission along the margin dominates over transmission to the surrounding tissue, as suggested by the patterns of cell shapes and tissue tension within the epiblast (*11*), the margin can be seen as a channel for this signal.

Our study, in which theoretical and experimental approaches intertwine, shows that it is the combination of both self-organizing and gene regulatory features of mechanical forces that enables embryonic regulation. On the one hand, self-organized tissue mechanics ensure that a single site of contraction forms in the intact epiblast, but also enable the rapid redirection of tissue motion (within hours if not minutes) in response to perturbations. On the other, by modulating gene expression, emergent tissue mechanics can refine embryonic territories to ensure their correct proportioning in intact embryos and trigger the emergence of ectopic embryonic axes in response to perturbations. Future work will be required to decipher how tissue mechanics elicits differential gene expression.

Finally, we note that although self-organized tissue mechanics can support spontaneous symmetry breaking, as manifested in ectopic embryo formation, it does not account for the original symmetry breaking event in normal development. Indeed, Gdf1, whose expression is restricted to the posterior epiblast before gastrulation movements initiate, is itself required for primitive streak morphogenesis and can redirect tissue motion when applied ectopically. Thus, tissue mechanics (at least, as we describe it here) functions as a canalizing feedback rather than as the initial trigger in normal axis formation. This illustrates the high level of interdependency between mechanical and molecular signals that safeguards development from deleterious deviations. Given the universal role of mechanical forces in shaping tissues, we anticipate that our findings and modeling framework will be relevant to patterning processes in diverse organisms, especially amniotes, including humans, which harbor a regulative and self-organized development.

## Materials and Methods

### Animals

All experimental methods and animal husbandry for transgenic quails were carried out in accordance with the guidelines of the European Union 2010/63/UE, approved by the Institute Pasteur ethics committee authorization #dha210003, and under the GMO agreement#2432.

### Embryo imaging, orientation, laser dissection, and drug treatment

Transgenic memGFP quail embryos (*11*) were collected at stage XI using a paper filter ring and cultured on a semi-solid nutritive medium of thin chicken albumen, agarose (0.2%), glucose, and NaCl, as described in (*25*). In addition, the nutritive medium was supplemented with H1152 dihydrochloride (Tocris, 25-75 µM) or Calyculin A (Tocris, 25 nM) for drug-treated embryos. The embryos were then transferred to a bottom glass six-well plate (Mattek inc.) with 2 mL of the nutritive medium (with or without drug) and imaged at 38°C with a microscope Zeiss LSM 900 using 2.5X or 5X objectives. The time interval between two consecutive frames was 6 min.

For bisection experiments, epiblasts were oriented by analyzing tissue flows in real-time using PIV. After 1-2 hours of image acquisition, the margin, the embryonic, extraembryonic territories and the presumptive anterior-posterior axis of the embryo were determined automatically, as previously described (*11*). Next, the spatial x-y coordinates of a line passing through the center of the epiblast, with a specific angle to the A-P axis (as explained in Fig. S2), were obtained and transferred to the ROE SysCon software of the laser dissector. Laser severing was performed using a UGA-42 firefly module coupled to a 355-nm pulsed laser (100% power) from Rapp Optoelectronic and the above-mentioned microscope and objectives. After bisection, the two halves of the embryo were moved on two separate vitelline membranes, or one half was left in its position and the other gently removed with a mouth pipette.

To ensure accurate comparison between posterior halves with fixed/free borders and intact epiblasts, their developmental stage was precisely matched. As described previously (*11*), embryos were staged according to the integrated contraction of a posterior segment of the margin. This was evaluated on the fly based on PIV analysis of tissue motion. Embryos exhibiting an identical contraction (20%, reached in ∼2 hours) were bisected and allowed to develop until they contracted by the same (45%, reached in ∼3 hours). The equivalent stage for control (intact) embryos was determined by calculating the total contraction (before + after the cut) of posterior halves (55%; ∼5h). Thus, embryos presenting the same amount of tissue converging towards the primitive streak are being compared, ruling-out possible confounding effect due to advection. For simplicity hours-only were reported in figures.

Attachment of the embryo was favored by cooling down the embryo at room temperature for 1 hour and removing excess liquid culture medium between the freshly cut edge and the vitelline membrane, whereas attachment was disfavored by imaging posterior halves immediately after the bisection. Embryos were subjected to a second laser dissection if the edges reattached, as in Fig. S4 and Movie S6.

### Beads Grafting and Obstacle

For the bead-based drug delivery experiments, Affi-Gel blue beads (100-200 µm diameter) were soaked at 4°C for four hours in HBSS solutions of 0.8% DMSO (0.8% v/v), Calyculin A (Tocris, 35 nM) and/or H1152 dihydrochloride (Tocris, 25-75 µM). The beads were then gently deposited, using fine forceps, in the anterior or posterior margin of either wild-type or memGFP embryos previously oriented by eye inspection. For the obstacle experiment, memGFP quail embryos were oriented by eye, and a fragment of hair (∼100 µm diameter, ∼6000 µm long) was gently deposited on the ventral side of the embryo.

### In Situ Hybridization

Quail embryos were fixed in ice-cold 4% formaldehyde, dehydrated in PBST (PBS/0,1%Tween) with increasing methanol concentrations (25%, 50%, 75% and 100%), and then rehydrated. Hybridization with DIG-labeled RNA probes was performed overnight at 65 °C in hybridization buffer (5X SSC pH4,5, 50% formamide, 1% SDS, 50 mg/mL yeast tRNA,50mg/mL heparin). The day after, the embryos were thoroughly washed and treated for six hours with a blocking solution of MABT, 2%BBR (Boehringer blocking reagent), 20%Lamb Serum. Then, the embryos were incubated overnight with an AP-coupled anti-DIG antibody in the blocking solution and finally stained with NBT/BCIP liquid substrate (Sigma). After the staining, the embryos were temporarily mounted on a slide and photographed at different magnification with a SteREO Discovery.V8, ZEISS (equipped with an AxioCam MRc, Zeiss). For the DIG-labeled probe, an 807 b.p. fragment of Snai2 coding sequence (from nucleotides 165 to 971) was PCR-amplified from quail cDNA and cloned in pGEM-T Easy vector (Promega).

### RNAscope

The samples were fixed overnight at 4 °C in ice-cold 4% formaldehyde and then washed in PBST (PBS/0,1%Tween) for six hours. The staining was performed using the RNAscope Multiplex Fluorescent Reagent Kit V2 according to the manufacturer’s instructions with the following modification for whole-mount staining of quail blastoderm embryos. For sample pre-treatment, no retrieval or proteinase treatment was performed. The probes used for the stainings are RNAscope Probe - Cja-GDF1 (Bio-Techne, 593421), RNAscope Probe - Cja-T-C2 (Bio-Techne, 587391-C2) and RNAscope Probe - Cja-LOC107312850-C3 (Bio-Techne, 587381-C3) and were detected with Opal520, Opal570 and Opal650 reagents (Perkin Elmer, 1/750 in TSA Buffer). Embryos were then mounted between slide and coverslip, using Fluoromount-G mounting medium (00-4958-02)

### Model

We considered a hierarchy of models to explore self-organization of force generation along the embryo margin through feedback of tissue tension on contractility. In the simplest instance, the margin is described as a 1D tensile line with a fixed length and periodic boundary conditions, and the surrounding tissue is ignored, corresponding to a limit where force transmission along the margin dominates over force transmission to the surrounding tissue. We can further take a limit where advection along the margin is negligible, which is the relevant regime for the embryo (regulation occurs on time scales that are shorter than the characteristic time scale for advection, which is the time scale over which the primitive streak emerges).

In the context of this minimal model, we identify contractility with the active tension *T*_*a*_ generated at the margin. Assuming that this active tension combines with a linear viscous resistance to stretching, the total tension *T*(*s*) at a point *s* along the margin is given by

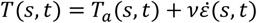

where 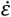 denotes the strain rate (the local elongation rate of the margin) and *ν* is a 1D viscosity. Assuming that the active tension varies in response to the strain rate, we write

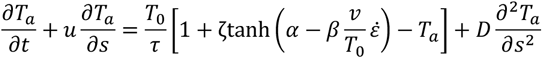

where *τ* is a characteristic time scale for regulation, and we have included a diffusion term with diffusivity *D* to account for non-local self-activation of contractility, such as might arise from the recruitment of neighboring cell-cell junctions into cables that span several cells, and are constantly turning over within the margin.

With the surrounding tissue being neglected, mechanical balance implies that the tension *T* is uniform along the margin, and conservation of the total margin length implies that the average strain rate vanishes, 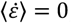, so that

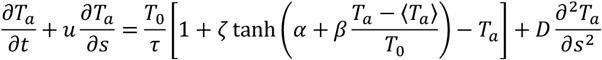

Thus, active tension is upregulated (resp. downregulated) where it is above (resp. below) its spatial average. Mathematically, this model is equivalent to a molecular activator-inhibitor model, where the tension *T* = ⟨*T*_*a*_⟩ serves as a long-range inhibitor, in the limit of infinitely fast and long-range inhibitor diffusion. If the coefficient *β* representing the strength of mechanical feedback is large enough, the model supports spontaneous symmetry breaking and the stable maintenance of regions of high and low contractility. In the relevant parameter regime, the steady state of the model is governed by the motion of narrow fronts between these regions, that tend towards invariant proportions, corresponding to a fixed total tension. When advection is taken into account, the fronts are displaced to a position where advection is balanced by the tendency to return to this preferred, ‘homeostatic’ tension. For details of the model and its analysis, see Supplementary Text.

Our full 2D model incorporates the same mechanical regulation of contractility into our previously described fluid-mechanical model of tissue flows in the embryo (*11*). Briefly, the embryonic disk is described as a 2D viscous fluid driven by tension along the margin, with a prescribed divergence term that allows for non-uniform areal expansion of the embryonic disk. The profile of stresses imparted on the tissue by the margin is assumed to keep an invariant, gaussian profile, such that distributed force generation at the margin and its regulation can still be described in terms of a 1D profile of contractility. To allow for a nonlinear relation between mechanical load on cell-cell junctions and contraction rate, and a saturation of the contraction rate, the margin is explicitly described as a 1D viscoelastic line; the rest length *l*_0_ of an element of margin varies as

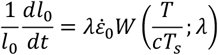

where *c* denotes the contractility (which is no longer identified with an active tension but could be understood to represent the local density of active myosin or supracellular cables) and *W* is a nonlinear, saturating ‘walking kernel’ (cf. (*26*)). *T*_*s*_ denotes a stall force per unit contractility at which junctions transition from contraction to yielding, and the parameter *λ* controls the nonlinearity of the walking kernel. With *E* denoting the elastic modulus of the margin, we obtain the system of equations

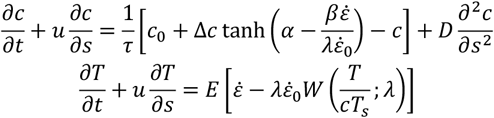

where *T* is the elastic tension along the margin, which determines the velocity *u* through the Stokes equation describing tissue motion. This 2D model is used to simulate the full course of margin regulation and tissue flows within the embryonic disk upon perturbations, and in a simplified geometry is amenable to a similar analytical description as the minimal 1D model. For numerical simulations, it was implemented in Python, using the FEniCS finite element platform. For a detailed discussion and analysis, see Supplementary Text.

## Supplementary figures

**Fig.S1.**
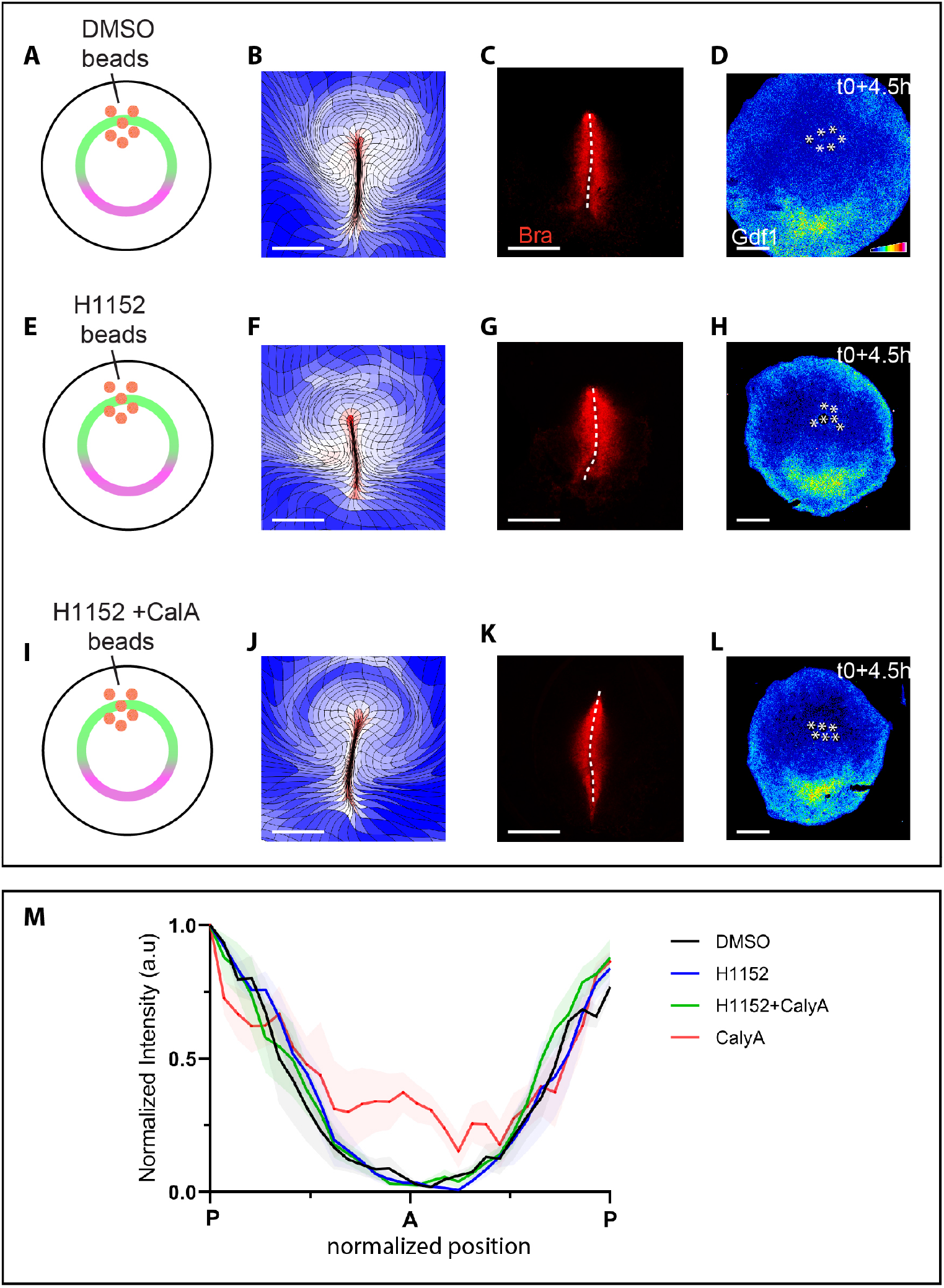
Control beads placed at the anterior margin do not trigger ectopic embryo formation. **(A to L)** Beads soaked in DMSO (A to D), H1152 (E to H) and H1152 and Calyculin A (I to L) and placed on the ventral side of unincubated quail embryos (asterisks denote the position of the beads). (A, E, I) Sketch of the experiment. (B, F, J) tissue deformation and expression of Brachyury (C, G, K) and Gdf1 (D, H, L) after bead deposition. (**M**) Normalized intensity profiles of Gdf1 expression at the margin and centered showing averages (colored curves) and standard error of the mean (shadowed region) embryos with beads soaked in DMSO (black, N= 4), H1152 (blue, N= 4), H1152 and Calyculin A (N= 4)or Calyculin A-only (N=4). Sacle bars, 1mm.

**Fig.S2.**
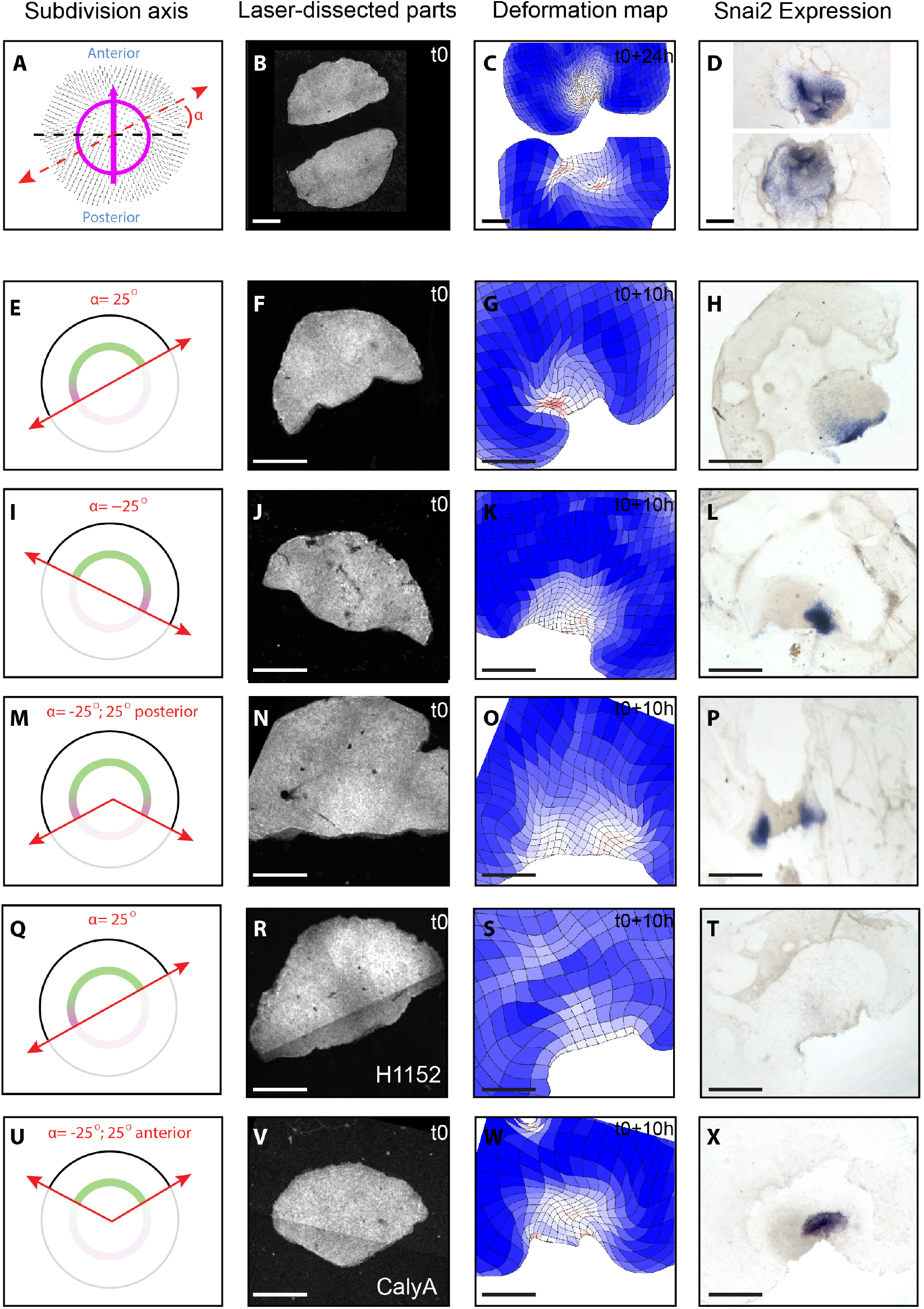
Formation of ectopic embryos in anterior parts bisected at varying angles. **(A to D)** Experimental procedure for the evaluation of bisection angles on ectopic embryo formation in anterior parts. Epiblast are bisected with an angle that is defined following the automated detection of the margin and anteroposterior axis using PIV analysis [(A); velocity fields, black arrows; margin and anteroposterior, in magenta; position of cut defined by the angle α, red dotted lines)]. Anterior epiblast parts are bisected using a UV-laser (B). The formation of ectopic axes is monitored by tissue deformation (C) and verified by (C) expression of Snai2. **(E to X)** Bisection experiments for different values of α (E, I, M, Q, U) showing anterior epiblast parts at t0 (F, J, N, R, V), the deformation maps after 10 hours (G, K, O, S, W) and the corresponding Snai2 expression. Scale bars, 1mm.

**Fig.S3.**
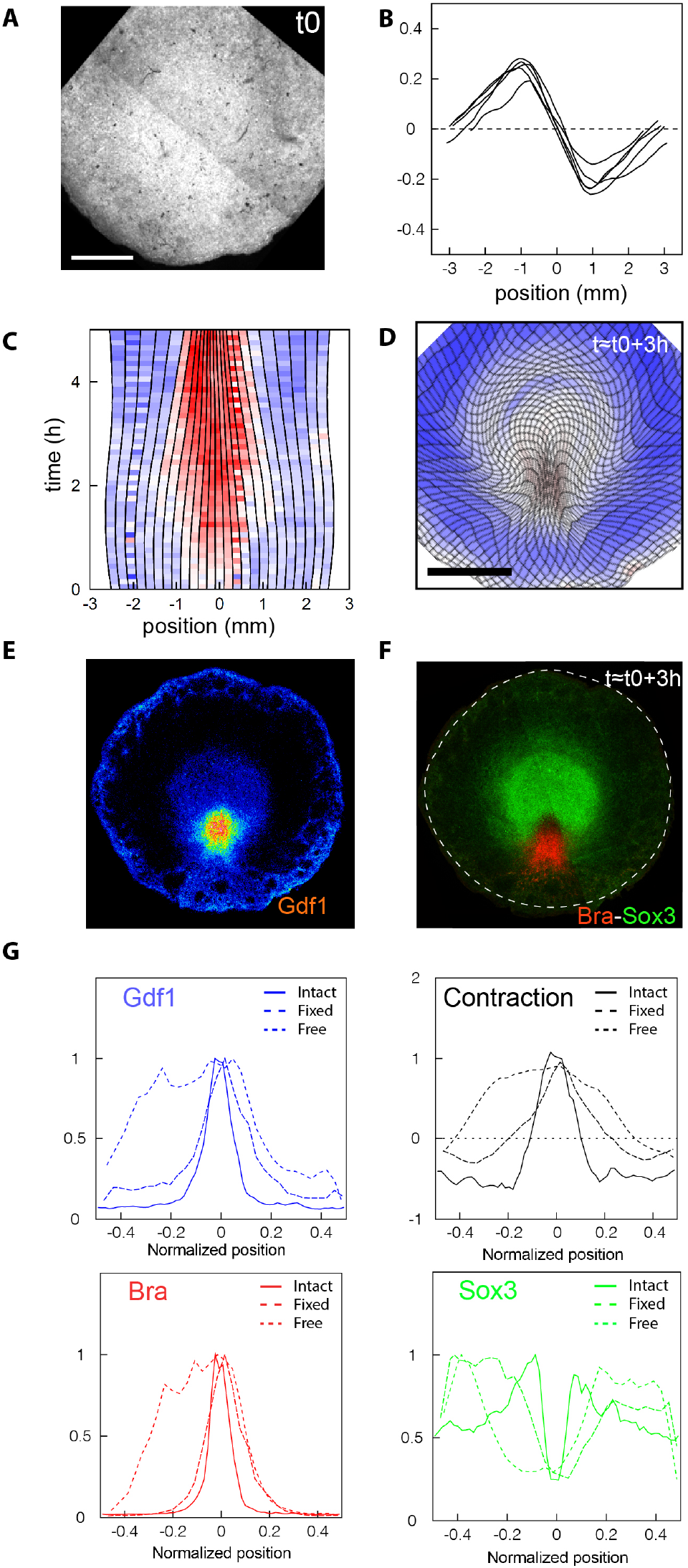
Tissue motion and embryonic territories in control intact epiblasts. **(A to F)** representative example of an intact epiblast whose developmental timing is precisely matched to posterior halves with free/attached boundary condition 3 hours after bisection (t=5h ≈ t0+3h, see Methods for details on the procedure). (A) Intact epiblast at t0. (B) Velocity profiles at the margin (N=5). (C) Elongation rates at the margin. Deformation maps (D) and corresponding expression of Gdf1 (E), Brachuyry and Sox3 (F) at t ≈ t0+3h (B to D). **(G)** Normalized intensity and contraction profiles at the margin averaged over N=3 embryos for each condition (free, attached and intact).

**Fig.S4.**
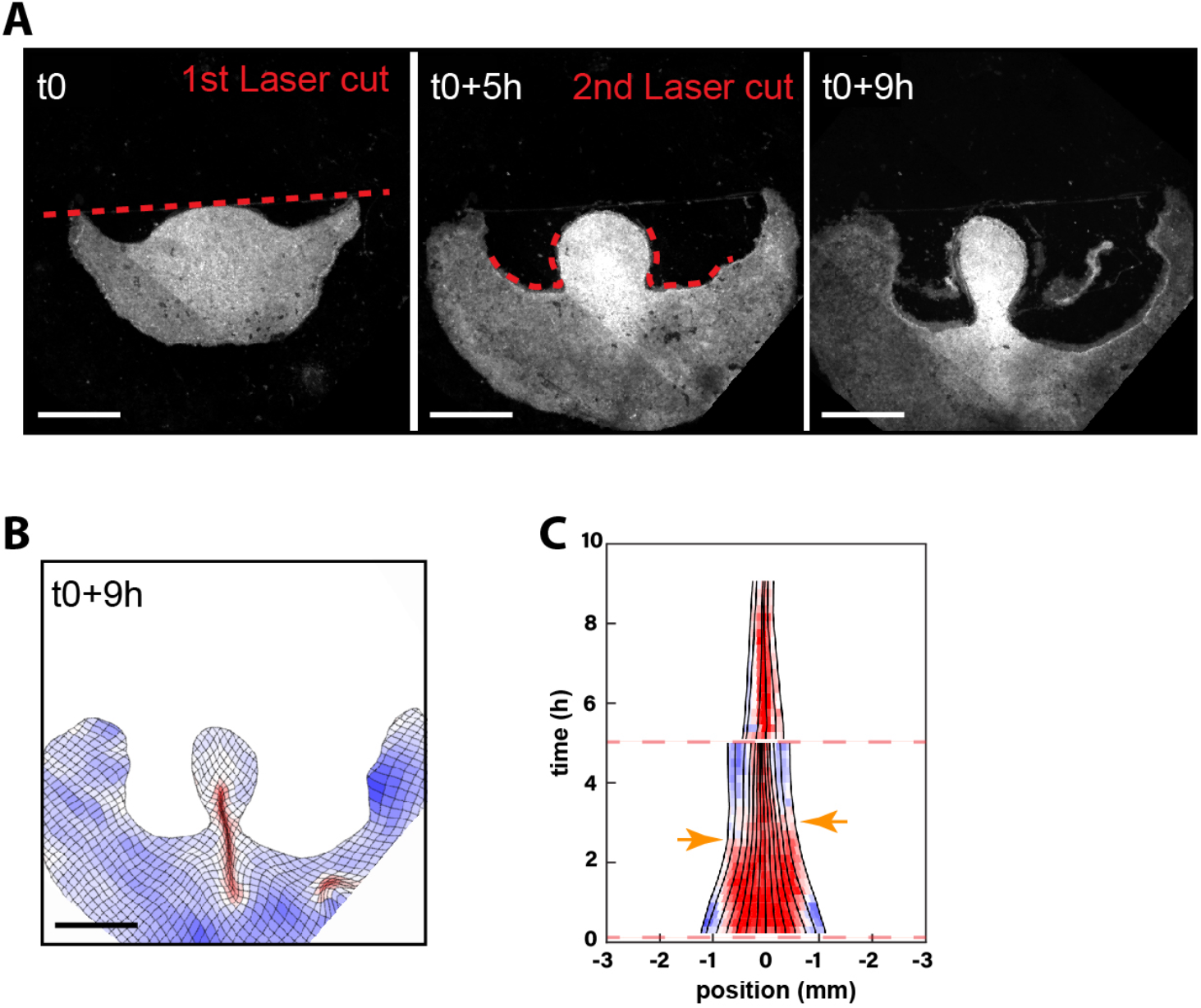
Long-term effect of posterior half with free edges. (A-C) Posterior half with repeated laser cuts to release border attachment. (A) time series of a posterior epiblast half just after the 1^st^ laser cut bisecting the epiblast (t0, left panel), just before the 2^nd^ laser cut, as embryo borders reattach (t0+5h, middle panel), and after the entire margin has converged and contributed to the forming primitive streak (t0+9hours, right panel). (B) Deformation map (t0+9h). (C) Corresponding kymograph of elongation rates at the margin (0mm is posterior; orange arrows point at the time epiblast borders reattach; red dotted lines correspond to successive laser cuts). Scale bars, 1mm.

## Supplementary movie legends

**Movie S1. A mechanical analog of Turing’s reaction-diffusion model for the regulation of tissue contractility**.

Left panel: Trajectories (the overlay denotes the embryo margin); velocity profiles at the margin and 2D deformation from an average of 6 different embryos between t0 and t8h;

Right panel: Mechanical self-organized model for the regulation of tissue contractility at the margin, with predicted contractility (magenta), tension (green) and velocity profiles (black) at the margin and 2D deformation.

**Movie S2. Tissue contractility modulates Gdf1 expression to drive the formation of a single embryo in intact epiblasts**.

Upper panel: Model predictions for localized perturbations of cell contractility showing profiles of contractility (magenta), tension (green) and velocity (black) at the margin and 2D deformation.

Lower panel: Experimental activation and inhibition of tissue contractility by deposition of drug-soaked beads showing original image acquisition and 2D deformation.

**Movie S3. Self-organized tissue mechanics drives ectopic embryo formation upon epiblast subdivision**.

Upper Panel: Model predictions for the response to epiblast bisection in anterior halves (upper panel) showing profiles of contractility (magenta), tension (green) and velocity (black) at the margin and 2D deformation.

Lower panel: Experimental bisections showing initial image acquisition, 2D deformation and corresponding expression of Gdf1 and Snai2 in halves +/- epiboly and treated with H1152.

**Movie S4. Procedure for monitoring embryo formation upon epiblast bisection**.

Intact embryos are imaged for 1-2h and analyzed using PIV on the fly. The orientation of the cut is defined following the automated detection of the margin and anteroposterior axis. The coordinates of the desired cut are transferred to the laser-dissection system. Bisected and separated epiblast halves are then fixed at early (∼5h) or late (>10h) time points and the expression of Gdf1 and Snai2 is then verified.

**Movie S5. Mechanical feedback (re)scales embryonic territories**.

Upper panel: Model predictions and experimental response to fixed and free boundary conditions in posterior epiblast halves after 3h. Contractility, tension and velocity profiles are in magenta, green and black, respectively.

Lower panel: Expression of Gdf1, Bra and Sox3.

**Movie S6. Long-term effect of mechanical boundary on embryonic territories in posterior halves**.

Left Panel: Posterior halve with attached borders, imaged for 8 hours.

Right panel: Posterior half with repeated laser cuts to release border attachment, imaged for 8 hours. Note the unbalanced contribution of the entire margin to the forming primitive streak at the expense of the epiblast territory.

**Movie S7. Mechanical friction redirects tissue motion, inducing ectopic embryo formation and rescaling of embryonic territories**.

Upper panel: Model predictions for localized friction across the epiblast showing contractility (magenta), tension (green) and velocity profiles (black) at the margin and 2D deformation.

Lower panel: memGFP embryo with a hair on its ventral side showing velocity profiles at the margin and 2D deformation maps at t0+14h and t0+4.5h with corresponding Gdf1, Bra, Sox3 expression.

## Supplementary Text

### I. MINIMAL ONE-DIMENSIONAL MODEL OF MARGIN REGULATION

#### A. Derivation and analogy with a Turing model

The goal of this model is to illustrate and analyze mechanical regulation at the embryo margin with a minimal number of parameters and variables. The fuller two-dimensional version that is employed to compare with experiments is presented in section II.

To begin, we consider the dynamics of the margin in isolation and ignore the influence of the surrounding tissue. Since the margin width is narrow compared to its curvature and length, we ignore any cross-sectional variations and consider a one-dimensional line, parameterized by arc length *s*. Along the margin we allow for tangential tissue motion with velocity *u*(*s, t*), which may also vary with time *t*, and define the contraction rate 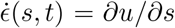. The contraction rate is positive when the margin is extending, and negative when it is contracting. We define *T* (*s, t*) to be the tension of the margin. This is composed of a passive contribution due to internal dissipation, which we take to be effectively viscous with a one-dimensional viscosity 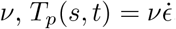, and a biologically active tension (or contractility) *T*_*a*_(*s, t*), which is mechanically regulated. We model this regulation with an advection-diffusion equation that includes a forcing term that depends on the extension rate 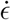,

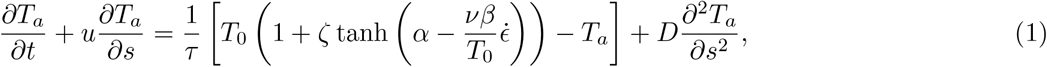

where *τ* is the time scale of the regulation, *T*_0_ a reference scale for the tension, *ζ* a dimensionless parameter indicating the amplitude of the regulation, *α* a dimensionless parameter controlling the crossover point in the response of active tension to stretching, *β* expresses the sensitivity of the regulation to stretching, and *D* is a diffusion coefficient that is added to phenomenologically incorporate a slightly non-local activation, as might be due to the recruitment of neighboring cell-cell junctions into cables that span several cells and undergo constant turnover within the margin. This equation describes the relaxation of the active tension to a steady state 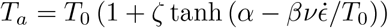. Fig. 1 shows an example of this steady state as a function of the extension rate 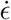 for parameters *ζ* = 1*/*3, *α* = −1, *β* = 3. The model hence captures the idea that (for positive *β*) active tension is downregulated in extending regions of the margin 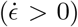 and upregulated in contracting regions 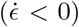, as could be due to tension-dependent breakdown of the cables.

**Figure 1:**
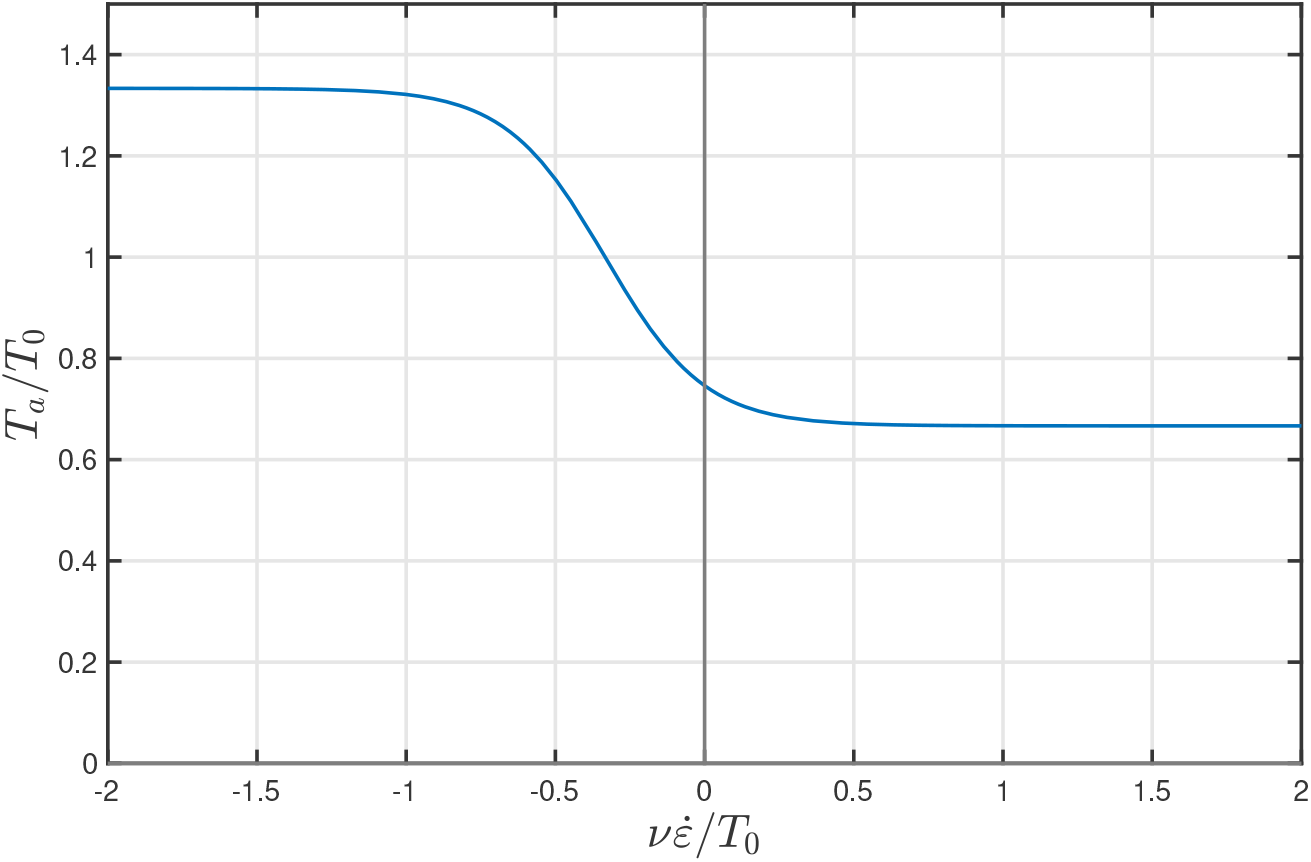
Steady state active tension *T*_*a*_ as function of the local margin extension rate 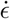 in for the minimal model, for the parameters *ζ* = 1*/*3, *α* = −1, *β* = 3. In extending regions 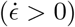 the active tension is downregulated to *T/T*_0_ = 1 − *ζ* while in contracting regions it is upregulated to *T/T*_0_ = 1 − *ζ*. The ratio *α/β* controls the location of the transition and the parameter *β* its width.

In order to close the minimal model, we need to describe additionally how the contraction rate responds to stresses in the tissue. From the definition of tension we have that

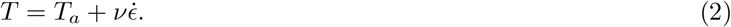

To proceed we exploit the fact that for the minimal model we are considering the margin in isolation from the tissue, which amounts to neglecting friction with the surrounding tissue compared to the internal stresses within the margin. Mechanical balance then implies that the total tension is uniform in space and varies only with time,

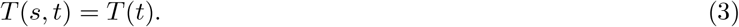

If the ends of the margin are closed in a loop (as in the case of the intact embryo), or fixed (as in the case of an ablated embryo where the cut has reattached to the vitelline membrane), the margin can sustain a non-zero tension *T >* 0. In contrast, when the ends are free (no reattachment) then *T* = 0. In the former case we can determine the value of *T* by considering the average of Eq. (2) over the length of the margin. Since the ends are fixed, so is the overall length and therefore the average contraction rate must be zero. Hence

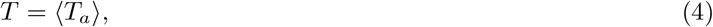

where angled brackets indicate a spatial average, 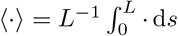, with *L* being the length of the margin. Thus, the uniform total tension is equal to the average of the active tension, and the contraction rate is given by

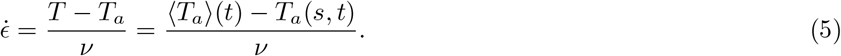

The tissue thus contracts (resp. extends) when the local value of the active stress is above (resp. below) its average value along the margin. Mechanical tension thus functions as a signal that inhibits contractility at a distance from the regions where it is highest, akin to the diffusing inhibitor in a molecular activator-inhibitor model, although on the long time scales of development, the transmission of mechanical forces is near instantaneous.

Combining Eqs. (1) and (5), we find that a single non-local equation governs the evolution of the active tension. We can gain further insight by scaling the variables to make all parameters dimensionless. To this end, time is scaled by *τ*, stresses by *T*_0_, and lengths by *L*, while the contraction rates are scaled by *T*_0_*/ν* and velocities by *T*_0_*L/ν*. The governing equation then reads

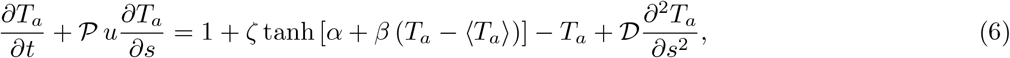

where two new dimensionless parameters are defined as

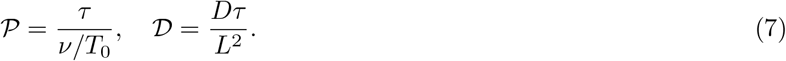

The dimensionless quantity 𝒫, which determines the importance of the advection term *u∂*_*s*_*T*_*a*_ in Eq. (6), is the ratio of the active tension regulation time scale *τ* to the time scale *ν/T*_0_ of advection of cables moving with the tissue. Since the experiments show that mechanical regulation occurs on the scale of minutes or a few dozens of minutes, while the tissue flows occur on the scale of hours, this number 𝒫 is small. Similarly, the effect of non-local activation is limited to scales much shorter than the overall length of the margin, and so 𝒟 ≪ 1. This leaves the minimal model with the governing equation (6) above and five dimensionless parameters, *α, β, ζ*, 𝒫 and 𝒟. In the following section we demonstrate that under the right conditions this model allows for the emergence of coexisting contractile and extensile regions.

#### B. Analysis of the minimal model

##### 1. Condition for the formation of contractile and extensile regions

In this section we demonstrate that the minimal model can predict the coexistence of steady contractile and extensile regions along the margin. We note that Eq. (6) allows for a trivial solution,

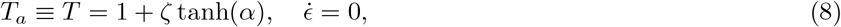

corresponding to a state with no motion and uniform tension *T* = 1 + *ζ* tanh(*α*). Spontaneous segregation into extensile and contractile regions occurs when this uniform state is unstable. In order to examine stability, we consider an infinitesimal perturbation *T*_*a*_ = *T* + *η* exp (2*πins* + *σ*_*n*_*t*) with *η* ≪ 1 and *n* ≥ 1 indicating the mode of the perturbation. Substitution into Eq. (6) yields

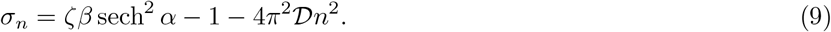

Hence the most unstable mode is the first, *n* = 1, corresponding to a single contraction, and the condition for instability is

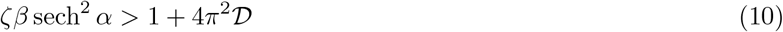

We conclude that for an instability to occur it is necessary that the regulation amplitude *ζ* and the regulation sensitivity *β* are sufficiently large. Strong biases in the regulation, corresponding to large absolute values of *α*, discourage an instability. Diffusion acts to suppress a short wavelength instability (below a length scale 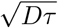), but a too great value suppresses the instability altogether. Since 0 *< ζ* sech^2^ *α <* 1, the condition requires in practice that the sensitivity *β* is quite large, and we shall frequently make the approximation *β* ≫ 1 in what follows.

The linear stability analysis presented above is limited to small perturbations about the uniform state, and so does not yield information on the behavior of the fully developed steady state with contractile and extensile domains, in particular it cannot account for the effect of advection and the motion of fronts. These are addressed in the next sections.

##### 2. Phenomenology of a bi-contractile state

In steady state *∂*_*t*_ = 0 and from Eq. (6) it follows that

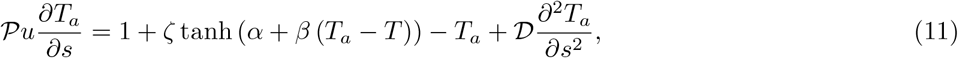

which we may rewrite in terms of the extension rate as

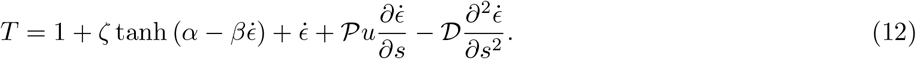

Here the left-hand side is constant along the margin, while the right-hand side depends on the extension rate. Thus, if multiple values of the extension rate 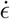 satisfying Eq. (12) exist, then the system allows for the formation of separate steady regions with different 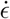. Away from fronts, the extension rate is approximately uniform and the advection and diffusion terms negligible. In this case

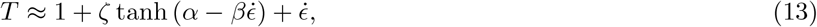

A ‘bi-contractile’ state with both contractile 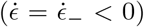 and extensile 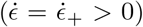 regions is guaranteed to exist if the instability criterion Eq. (10) is satisfied, and the system converges to it on a time scale ∼ *τ*, though due to the slow motion of fronts between the regions it may take longer to reach a steady ratio of the two domains. This is illustrated in Fig. 2. In addition, we present an example of the steady state as predicted by the minimal model in Fig. 3.

**Figure 2:**
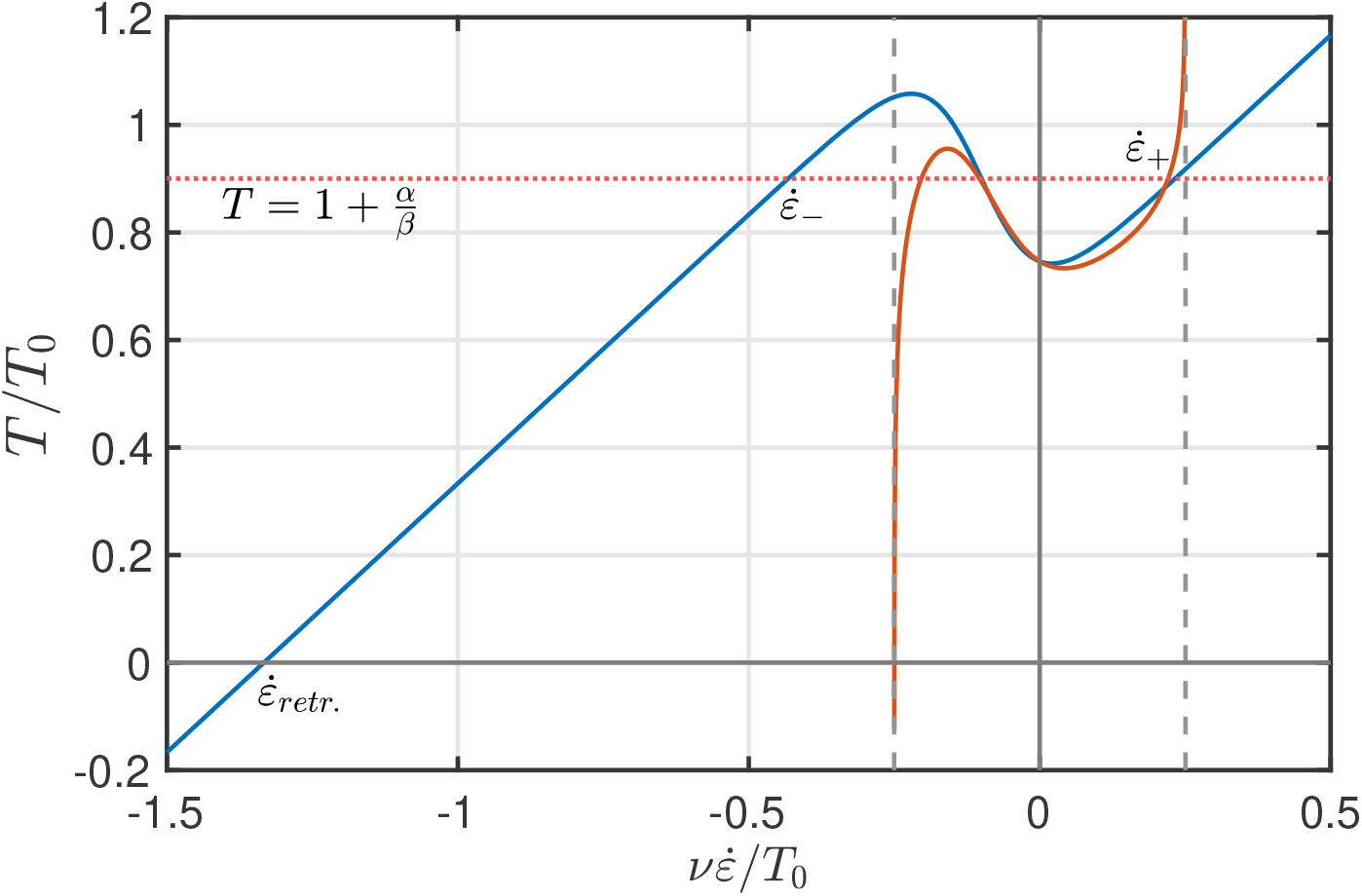
Illustrative steady state total tension for the minimal model (blue) and the full model model with non-linear walking kernel (orange). For the minimal model, the steady state tension can be calculated approximately (ignoring corrections from advection and diffusion), predicting the coexistence of extensile 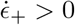 and contractile 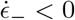 regions. In the absence of tension, as in the case of ablation, the model predicts a uniform negative contraction rate 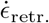. While the minimal model predicts a large ratio 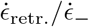, this is rectified through the introduction of the non-linear walking kernel in the full model. Parameters: *α* = −1, *β* = 10, *ζ* = 1*/*3, *λ* = 4.

In order to determine the stationary size of the contractile domain, which we call *ρ*, it is helpful to consider an analogy of Eq. (13) with a ball rolling in a potential with friction [1]. In the limit where tissue advection is negligible, 𝒫 → 0, there is no friction and fronts in the system may be understood as a transition between maxima with equal potential. By symmetry of the regulation, it follows that the inflection point of the regulation, 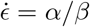 is a stationary point (local minimum) as well, 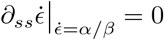. Hence

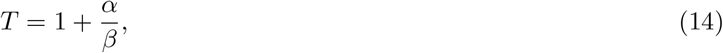

in a steady bi-contractile state. For sensitive regulation, *β* ≫ 1, this implies

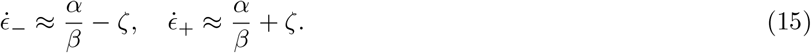

**Figure 3:**
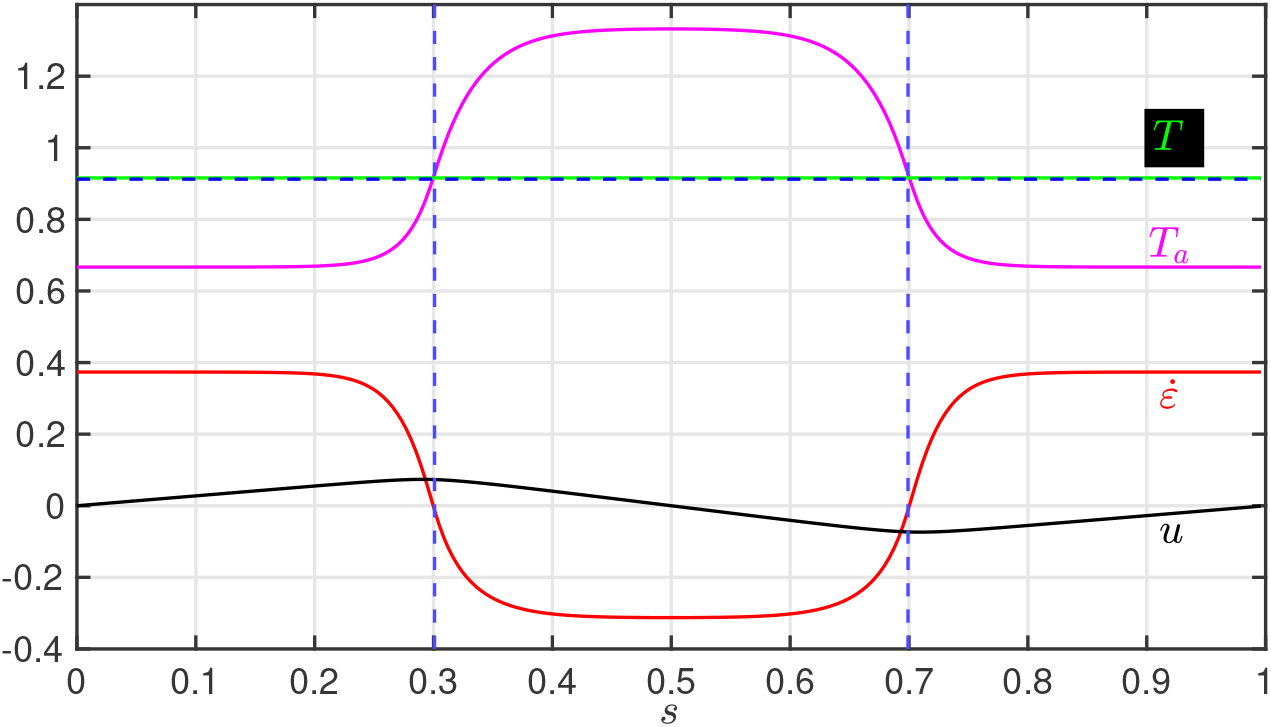
Illustration of the steady state predicted by the minimal model. The vertical axis displays tension *T* (green), active tension *T*_*a*_ (magenta), extension rate 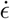 (red) and velocity *u* (black). All quantities are scaled to be dimensionless. Blue dashed lines indicate theoretical predictions for the size of the contractile domain *ρ*, Eq. (25), and the tension *T*, Eq. (24), using the value *u*(*s*_*f*_) = 0.0737 obtained from the simulation to determine the Péclet number Pe.. Simulation parameters: *α* = 0, *β* = 16, *ζ* = 0.33, 𝒟 = 7.5 × 10^−4^, 𝒫 = 0.2.

Taking the spatial average of Eq. (12) we also find that

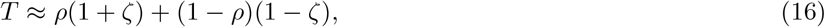

where we assumed that the fronts are narrow compared to the size of the domain and used that 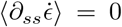 by periodicity. In conclusion,

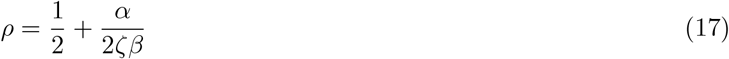

in the limit 𝒫 → 0 (regulation is fast compared to advection). From this we see clearly that the parameter *α* controls the size of the contractile domain, with negative *α* favoring a more localized contraction, as observed in experiments. In particular, for *α* = 0 this simple calculation invariably predicts *T* = 1 and *ρ* = 1*/*2. Comparing with Fig. 3 we see that these predictions are slightly too large and there are corrections when the advection 𝒫 is non-zero. These may be obtained by analyzing the behavior at fronts more carefully, as shown next.

##### 3. Fronts and the approach to steady state

In order to gain further physical insight into the steady non-linear state, and in particular to understand the role of tissue advection, it is useful to consider first the nature of the contractility profile at the fronts between extensile and contractile regions. Without loss of generality we consider a front at *s* = 0 with extension for *s <* 0 and contraction for *s >* 0. The centerpoint of the front is defined to be the inflection point of the regulation term, i.e. such that 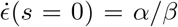. Assuming a high sensitivity, *β* ≫ 1, we may treat the regulation as approximately bimodal, with *T*_*a*_ → 1 − *ζ* in the extensile and *T*_*a*_ → 1+*ζ* in the contractile region. Eq. (6) then reduces to the steady one-dimensional advection-diffusion equation with piecewise constant forcing,

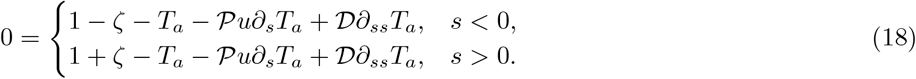

Assuming that the fronts are narrow compared to the length of the domain (consistent with velocity profiles in experiments), the velocity *u* varies only slightly across the width of the front and we can approximate its value by a constant *u* ≈ *u*_0_. The solution for the active tension is then

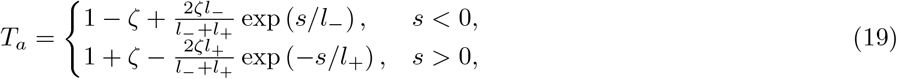

where

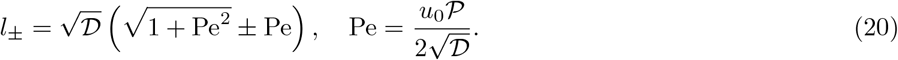

Here we define a Péclet number Pe that measures the relative importance of advection and diffusion in shaping the front. The velocity *u*_0_ may be estimated from the behavior of the model in the absence of advection. Since the extension rate in the contractile region is approximately constant, 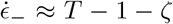, the profile of *u* is approximately triangular, and

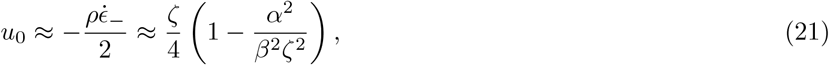

where we used the approximations for 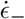 and *ρ* in the case without advection, Eqs. (15) and (17). It follows that the Péclet number can be approximated as

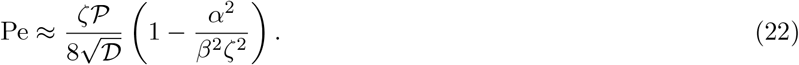

We see that passive regulation along the margin forms the profile of the front into different shapes upstream and downstream. The upstream length *l*_−_ is compressed, while the downstream length *l*_+_ is extended when 𝒫 *>* 0. The overall dimensionless width of the front *l*_+_ + *l*_−_ scales as 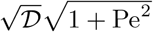 and is thus controlled by a combination of propagation and diffusion, and the assumption of a small front width is justified if 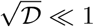 and Pe is not too large.

The time scale of advection by tissue flow is *ν/T*_0_, i.e. the time scale of mechanical tissue contraction. Experiments suggest that typical contraction rates are on the order of 0.2 hr^−1^, corresponding to a tissue motion time scale of about 5 hours, which is much longer than the biological regulation which is supposed to occur on the scale of minutes or a few dozen minutes. As a result, the model generically predicts the formation of a bi-contractile state with moving fronts that gradually converge towards steady state. The velocity of such a front *v* may be approximated by considering a solution *T*_*a*_(*s*_*f*_ = *s* − *vt*) to Eq. (6) and integrating it across a front after multiplication by *∂*_*s*_*T*_*a*_. From Eq. (19) we have 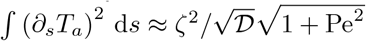 and so in the limit of sensitive regulation *β* ≫ 1 we find

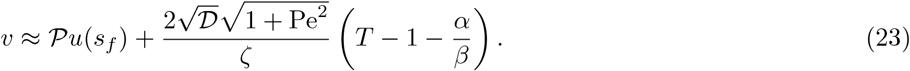

From this we learn that the velocity at which the fronts move relative to the tissue is controlled predominantly by the tension in the margin, with lower tension promoting the expansion of the contracting domain. In steady state *v* = 0 and refined approximations for the tension and domain size are found to be

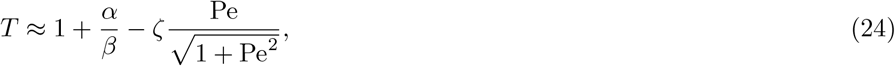

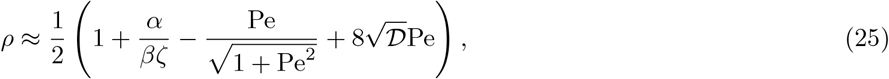

where the Péclet number is defined in terms of the velocity at the front, 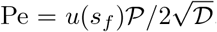. Comparing again with Fig. 3, we see that these estimates are significantly more accurate. To be exact, the error in the estimate for *T* is 0.3%, and in the estimate for *ρ* it is 0.6% when the Péclet number is calculated using *u*(*s*_*f*_) taken from the simulation, and about 1.4% and 3.0% when Pe is approximated using Eq. (22). Conversely, taking the values of *T* and *u*(*s*_*f*_) from the simulation we find that Eq. (23) predicts *v/u*(*s*_*f*_) ≈ 0.5%.

In conclusion, convergence towards a steady pattern in the model results from the motion of fronts between the contracting and stretched domains, which itself depends on the balance between advection, which tends to narrow contracting domains, and a ‘restoring force’ that tends to bring the proportions of the domains to a point where tension along the margin matches a ‘homeostatic tension’ 1 + *α/β*. The strength of this restoring force depends on the effective diffusivity, i.e. on the nonlocal activation of contractility, and on the rate of regulation, which must both be large enough to maintain a stable contractile domain.

##### 4. Release of tension and formation of uniform contractile domain

The analysis so far has assumed a strictly positive tension, as in the uniform steady state. This explains the regulation of contractile and extensile regions in a periodic domain, or a domain with fixed end points as in the case of reattachment after ablation. In particular, since the ratio of contractile and extensile domains *ρ* depends only on *α* and not on the length of the margin, the model predicts that the contractile region rescales after ablation if the ends reattach (provided the instability criterion Eq. (10) is still satisfied, but this also depends only weakly on the size of the domain through the non-dimensional diffusivity 𝒟, and so is in general still going to be satisfied for reasons outlined previously). In contrast, when the ends do not reattach, the model predicts very different behavior. In this case the overall tension is rapidly reduced to *T* = 0. As illustrated in Fig. 2, this tension generically lies outside of the admissible window for bicontractility and there is only a single stable steady state, corresponding to a uniform contraction with

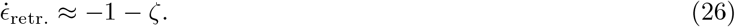

While this is in agreement with experimental observations qualitatively, we note that the ratio between the release contraction rate and the contraction rate in steady state is predicted to be rather large,

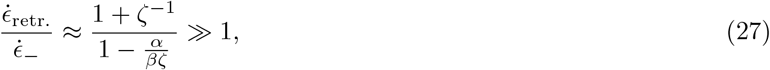

for reasonable values of the parameters. This contrast with experimental observations is ultimately due to the limitations of the minimal model. This is rectified by means of a non-linear description of the contractility dynamics in the full model in section II A and also illustrated in Fig. 2 (orange curve).

### II. TWO-DIMENSIONAL MODEL WITH TISSUE FLOW AND NON-LINEAR REGULATION

#### A. Overview

While the minimal model serves well in providing a qualitative understanding of the self-regulation, it does not provide for a complete picture. In order to address its shortcomings, we now introduce a full two-dimensional model of the regulation dynamics in the embryo that differs in two key aspects: first, the regulation dynamics of the margin are fully coupled to the motion of the underlying tissue, and second the generation of active stresses in the actomyosin cables that constitute the margin is modeled explicitly in terms of myosin motors. By modeling the tissue motion and growth we are able to apply the model directly to different embryonic geometries and boundary conditions, such as ablated posterior and anterior halves, and numerically predict the location and size of contractile regions, as well as the morphology of the embryo. By allowing for a nonlinear relation between the load borne by supracellular cables and the rate at which they contract, the extended model is able to capture the controlled contraction speed after ablation. It also takes into account explicitly the three different time scales of the dynamics, namely those for biological regulation, internal force transmission along the actomyosin cables, and advection speed of the tissue, as opposed to the minimal model which focused on the biological regulation only.

Departing from the minimal model that represents margin contractility in terms of a time-dependent active tension profile, we now explicitly consider the actomyosin cables that make up the margin as active, viscoelastic elements, with a nonlinear behavior that includes a saturation of the contraction rate in the case of very high or low tension. Specifically, the rest length *l*_0_(*s, t*) of a margin element responds a given tension *T* (*s, t*) as

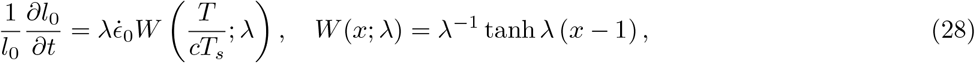

where define the walking kernel *W* in reference to [2]. Here we introduce *c*(*s, t*) as a variable for contractility, which can be understood to represent the local density of active myosin or supracellular cables, and *T*_*s*_ a stall force per unit contractility at which junctions transition from contraction to yielding. As is illustrated in Fig. 2, the kernel *W* is constructed in such a fashion to allow for viscous sliding when the load *T* is close to the equilibrium value *cT*_*s*_, but saturates to a maximal contraction rate, 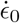 tanh *λ*, when the tension is released, *T* = 0. Here *λ* regulates the degree of non-linearity and for *λ >* 1 the maximal contraction rate is very close to 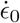. The kernel captures the idea that in the absence of tension myosin walkers contract with a fixed rate 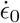 that is independent of the number of walkers present. If the load is increased, the walkers need to overcome resistance to motion and stall at a load of *T*_*s*_ per unit concentration. When the load is increased further, the actomyosin cables begin to yield and eventually reach a maximal extension rate. An illustration of the walking kernel is provided in Fig. 4.

Having described the time evolution of the rest length of individual margin segments, we can write an equation for the evolution of elastic tension along the margin,

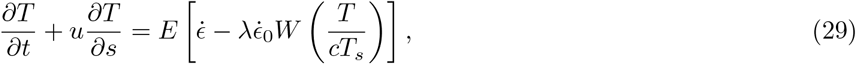

where *E* is the elastic modulus of the actomyosin cables.

**Figure 4:**
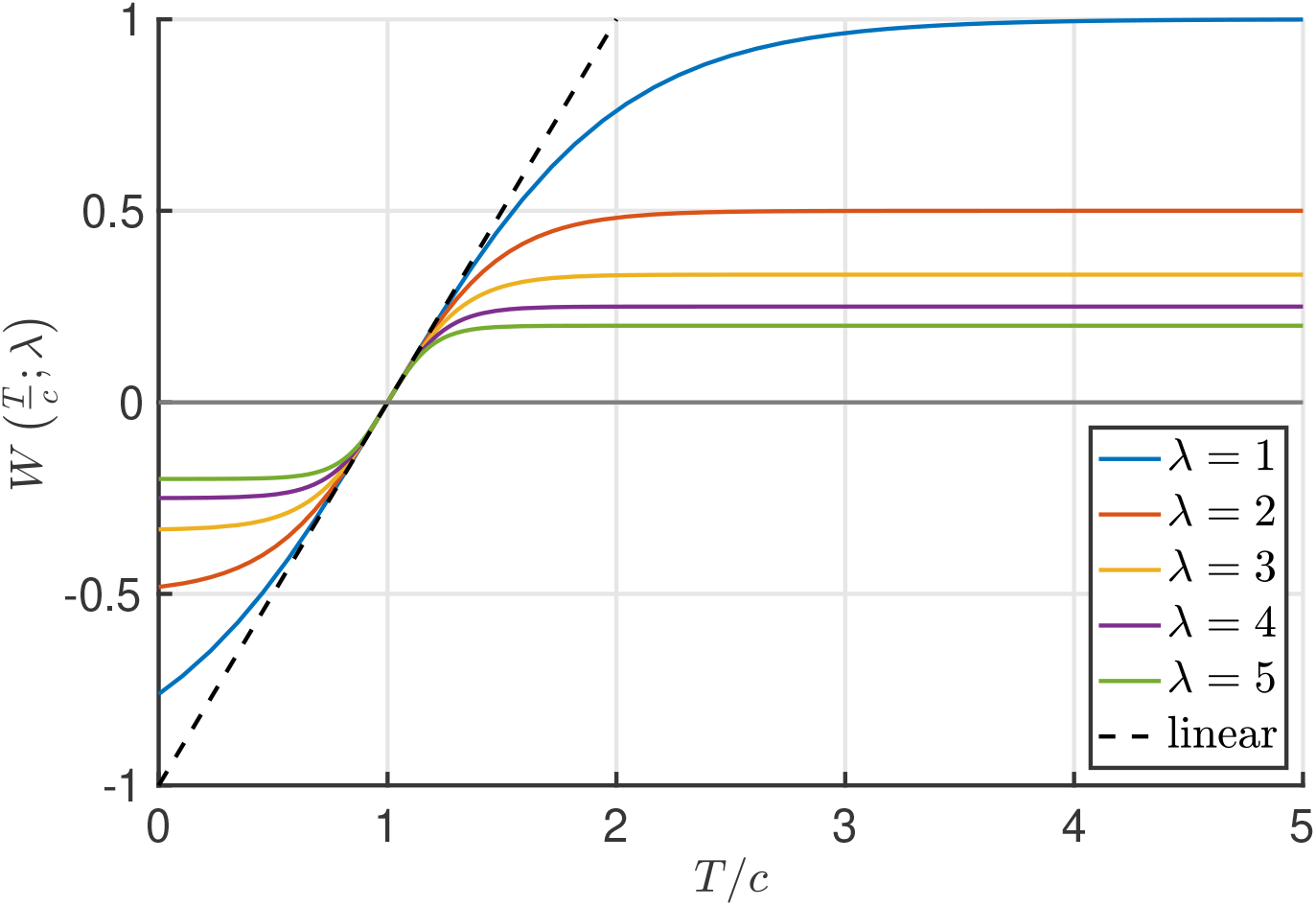
Illustration of the walking kernel *W* (*T/c*; *λ*) for different values of *λ* and comparison with the linearised kernel Eq. (41) for the case *ζ* ≪ 1. The parameter *λ* controls the degree of non-linearity. In the absence of tension, *T* = 0, the walking kernel is bounded by −1*/λ*, ensuring that contraction rates are not too big when tension is released.

With force generation now explicitly described in terms of active, viscoelastic elements, we transpose the equation for the regulation of active tension in the minimal model, Eq. (1), as an equation for the contractility *c* or density of these active elements,

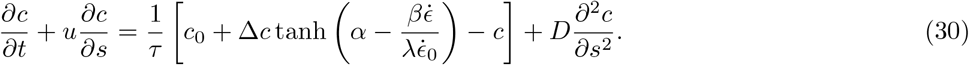

Here we introduce a reference contractility *c*_0_ and contractility amplitude Δ*c*. To incorporate the coupling between the margin and the surrounding tissue, we follow [3] and model the embryonic disk as a two-dimensional Stokes fluid with a prescribed divergence *γ*(***x***, *t*) and active stresses ***σ***_*a*_(***x***, *t*),

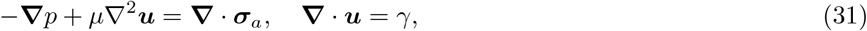

where *µ* is the dynamic viscosity of the tissue, ***u***(***x***, *t*) the 2D flow velocity and *p*(***x***, *t*) the pressure. Treating the contractile cables within the margin as viscoelastic but the surrounding tissue as a fluid greatly simplifies the analysis of motion driven by the margin, and is also justified by the observation that tissue outside the margin more readily dissipates tension, whereas larger elastic strains build up in the margin, as revealed by laser ablation experiments [3].

The margin is immersed in the tissue and the locus of its centerline is described by the position vector ***r***(*s, t*); by definition of *s* as arc length the tangent ***t*** := *∂****r****/∂s* is a unit vector. The active stress in the tissue ***σ***_*a*_(***x***, *t*) is a second-rank tensor and instantaneously given by

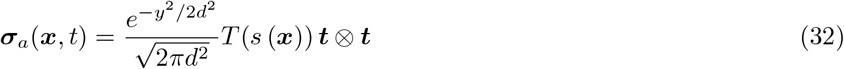

where *s*(***x***) corresponds to the point on the margin closest to ***x***, *y* = |***x*** − ***r***(*s*(***x***)) | and *d* a parameter that provides a scale for the margin width. In the limit where the margin width *d* → 0, the force distribution becomes singular on ***r***(*s*). This model captures the idea that regulation along the margin occurs without any cross-sectional variation, and hence remain a one-dimensional problem described by Eqs. (29) and (30) with *u* = ***t*** · ***u***(***r***) and 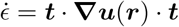. At the same time, the stress profile across the margin is described by a Gaussian to reflect the fact that it is slightly fuzzy with multiple actomyosin cables spread in parallel. The normalization is chosen to ensure that the net stress applied to the tissue is independent of the width parameter *d*.

In summary, the regulation dynamics are fully described by the coupled system Eqs. (29)-(32). Conceptually, the instantaneous distribution of stresses in the margin determines the tissue flow through Eq. (31), which in turn informs the evolution of biological contractility and mechanical stresses in Eqs. (29)-(30) and the advection of margin elements with the tissue. This extended model exhibits a number of qualitative differences compared to the 1D minimal model, the most significant of which are a bound on the contraction rate and the possibility (indeed, the requirement) of a non-uniform tension profile *T*. In the following sections, we analyze these differences in detail and explain why they are necessary to accurately describe the biological experiments.

#### B. Analytical simplification in a straight periodic geometry

The analysis of the complete model is greatly simplified when the margin is assumed straight, with ***t*** constant, since this allows for an explicit solution of the Stokes equations Eq. (31). For the theoretical analysis we also ignore the effect of area changes, *γ* = 0, since these do not directly influence the tension profile along the margin.

We imagine a Cartesian coordinate system (*x, y*) with the margin centerline on the *x*-axis and period *L*. The tissue velocity ***u*** decays to zero as *y* → ± ∞, far away from the margin. Ignoring the effect of tissue growth, the flow field is incompressible and hence can be described by a scalar stream function *ψ*(*x, y*) that satisfies the biharmonic equation ∇^4^*ψ* = 0. The velocity field may then be recovered as ***u*** = (*u*_*x*_, *u*_*y*_) = (*∂*_*y*_*ψ*, − *∂*_*x*_*ψ*).

Because of the periodicity of the margin, the tension may be written in terms of Fourier modes 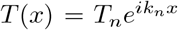 where *k*_*n*_ = 2*πn/L* and *n* is a positive integer labelling the mode. Using an ansatz with similar periodicity for the stream function, 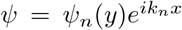, we find that the Stokes equations Eq. (31) simplify to an ordinary differential equation for *ψ*_*n*_,

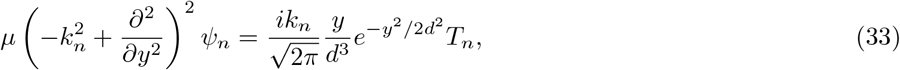

which can be solved to find *ψ*_*n*_ explicitly in terms of exponentials and the error function. Requiring boundedness as *y* → ± ∞ and up-down symmetry is sufficient to ensure uniqueness of the solution. From this, it may be shown that the *n*th mode of the horizontal extension rate, defined as 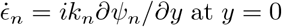 at *y* = 0, is related to the *n*th tension mode *T*_*n*_ by

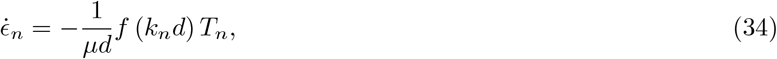

where the transfer function *f* is given by

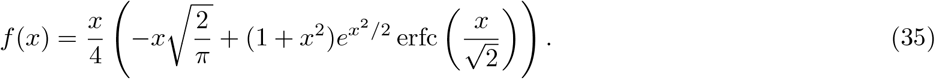

The response function *f* is strictly positive, grows linearly for long wavelengths (*f* ∼ *k*_*n*_*d* as *k*_*n*_*d* → 0) and decays quadratically as *k*_*n*_*d* → ∞. The decay for large *k*_*n*_*d* ensures mechanical robustness of the tissue flow to small scale perturbations. It is due to the non-local entrainment of the tissue by the Gaussian distribution of actomyosin cables, and disappears in the limit of zero margin width, *d* → 0.

Further insight may be gained by scaling the model equations to obtain all dimensionless parameters. To this end, we scale contractility by *c*_0_, tension by *c*_0_*T*_*s*_, lengths by *L*, time by the regulation time scale *τ*, while the extension rate is scaled by 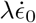 and velocity by 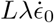 for reasons that will become clear shortly. This leads to the three scaled equations

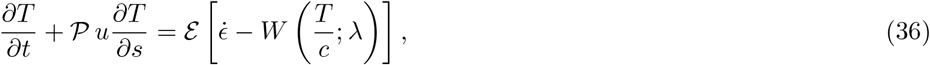

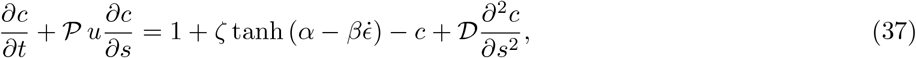

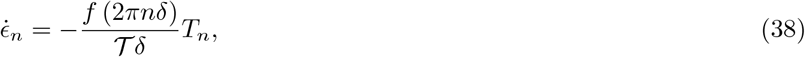

where we define

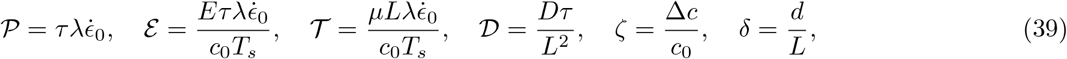

which together with *α, β* and *λ* brings the number of model parameters up to nine, compared to five in the minimal model. While at first these dimensionless parameters may appear counter-intuitive, their meaning becomes clearer when we define 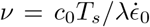 and *T*_0_ = *c*_0_*T*_*s*_ in analogy with the viscous internal dissipation of the margin in the minimal model. In this case several of the coefficients may be rewritten as

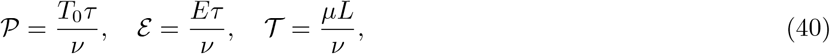

which not only makes explicit their connection with the minimal model, but also gives a physical meaning to ℰ as the ratio of elastic to viscous dissipation in the margin, and 𝒯 as the ratio of viscous dissipation in the tissue to that in the margin, in addition to 𝒫 that quantifies advection within the margin. Furthermore, in steady state the (dimensionless) contraction rate 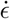 is bounded by *λ*^−1^, which makes it possible to interpret *λ* as a measure of the non-linearity of the walking kernel.

#### C. Reduction to the minimal model

It is now straightforward to recover the minimal model from the scaled equations of the complete model, Eqs. (36)-(38). Specifically, it emerges in the case when the actomyosin cables of the margin are very stiff compared to the viscous forces that are generated internally by the sliding myosin crawlers, and if both of them are much bigger than the viscous dissipation in the surrounding tissue. Mathematically, this corresponds to the condition *Eτ* ≫ *ν* ≫ *µL*, or ℰ ≫ 1 ≫ 𝒯 in terms of the dimensionless parameters from the previous section. In addition, it is necessary consider a margin of vanishing cross-section *δ* → 1, consistent with the picture of a 1D line in the context of the minimal model.

We demonstrate this by considering a small perturbation from the equilibrium solution of Eqs. (36)-(38). In the absence of any motion, 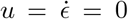 and the tension and contractility have the identical uniform value *T* = *c* = 1 + *ζ* tanh *α*. Considering the ansatz *T* = 1 + *ζ* tanh *α* + *T*′ and similar for *c*, we find that the walking kernel *W* may be linearized as

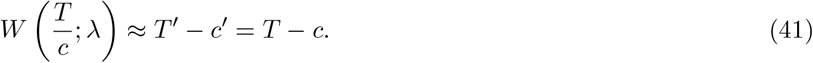

The condition ℰ ≫ 1 applied to Eq. (36) implies that the right-hand side must balance itself, and so

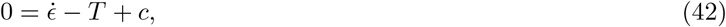

or, reinserting dimensions,

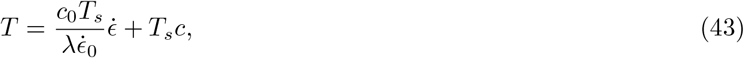

which is equivalent to Eq. (2) if we identify *T*_*a*_ = *T*_*s*_*c*. Hence we recover that the active tension in the minimal model is equivalent to the contractility in the linearized complete model. At the same time, this provides a justification for the definition of *ν* in the previous section as an effective linear viscosity that emerges through the sliding of the myosin motors. It is immediate then that Eq. (37) corresponds precisely to Eq. (1). Finally, for an infinitely thin margin *δ* → 0 we can Taylor-expand the function *f* to see that the right hand side of Eq. (38) is independent of *δ*. The condition 𝒯 ≫ 1 then guarantees that an 𝒪(1) contraction or extension is compatible with a small tension perturbation as assumed initially. The limitation of the argument by the approximation Eq. (41) restricting the margin tension *T* to small perturbations from its highlights clearly why the minimal model cannot capture the full behavior of the nonlinear model, as in the scenario where the tension is suddenly released.

#### D. Front motion in the nonlinear model

In a similar fashion to the minimal model, it is possible to gain insight into the motion of fronts in the extended non-linear model. Assuming that advection of tension (unlike advection of contractility) plays a limited role, as supported by our numerical simulations, we have 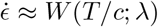. If we approximate the tension, like the velocity, as uniform across the front, *T* ≈ *T* (*s*_*f*_), we can rewrite Eq. (37) as

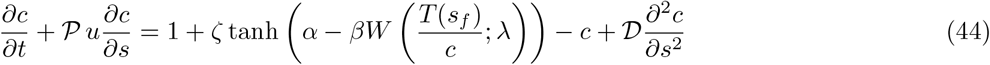

In the limit where *β/λ* ≫ 1, we can make the further approximation

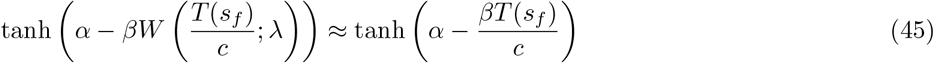

Thus, we recover an equation that is very similar in form to the equation for the time evolution of active tension in the minimal model Eq. (6). Following the same steps as in that case, we find that the same equation relating the velocity of the front to the local tissue velocity and tension, Eq. (23), applies. In order words, whereas a more extensive analysis of the full nonlinear model may be complicated by the nonlinear equation of motion relating tension, velocity, and contractility, its behavior can be understood in the same way, as resulting from a balance between advection and tension homeostasis.

#### E. Stability analysis and condition for the formation of a single embryo

In this section we consider the dimensionless system of governing equations Eqs. (36)-(38) of the extended model. We examine the stability of the uniform state *T* = *c* = 1 + *ζ* tanh *α*, 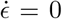 by assuming a perturbation of the *n*th mode, proportional to exp (*i*2*πns* + *σ*_*n*_*t*). Linearising the system then sets a quadratic condition on the growth rate *σ*_*n*_,

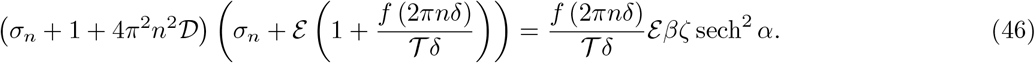

This enables a positive growth rate (and thus instability) if and only if

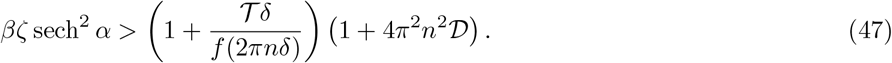

As in the case of the minimal model, which we recover in the limit of vanishing tissue viscosity 𝒯, Eq. (10), this essentially amounts to a condition on the regulative sensitivity *β* to be sufficiently large, while the spatial extent of actomyosin cables both tangentially (through 𝒟) and orthogonally (through *δ*, c.f. the asymptotic behavior of *f* discussed in section II B) to the margin ensures that short wavelength instabilities, as for example due to noise, are suppressed. In contrast however, the fundamental mode *n* = 1 is no longer guaranteed to be the most unstable, instead there is a competition of hydrodynamic and regulative effects.

In order to better understand the effect of the inclusion of tissue motion through the parameter 𝒯, it is instructive to consider the mode relationships between contractility, tension and extension rate. In the case of a stiff margin ℰ ≫ 1, we also have that *σ* ∼*βζ* sech^2^ *α* −1 and hence typically *σ* ≪ ℰ, so we recover approximately the linear relationship between the three quantities as in the previous section, 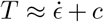. Using the long-wavelength approximation for *f* we then find that for a step-like profile of the contractility we have the approximate scalings

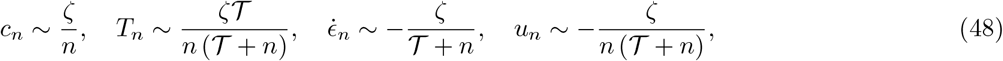

for *n >* 1. From this we see again that in the case of vanishing tissue viscosity 𝒯, there is no variation in tension. However, it also becomes evident that the inclusion of the tissue regularizes the contraction and velocity profiles 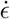 and *u* in response to the contractility *c* by acting as a high-pass filter. Thus the effect of the tissue manifests in a reduction of the magnitude of the extension rate in the center of extensile regions, as well as the generation of a tension peak in contractile regions.

Physically, the ‘drag force’ of the surrounding tissue screens the propagation of tension along the margin. If transmission along the margin is too strongly hindered, and the stretching imposed in extensile regions too strongly reduced, these regions may become unstable toward the formation of an ectopic contraction. This is reflected in the linear stability analysis carried out above, which shows that the fundamental mode *n* = 1 may not be the most unstable if the tissue viscosity 𝒯 is large, and is exacerbated in the non-linear regime where advection further shrinks contractile regions and extends extensile regions. We may illustrate this by considering a generalization of Eq. (47) to a perturbation about a state with a uniform extension rate 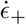 (as is approximately the case in the anterior). In that case an instability occurs if

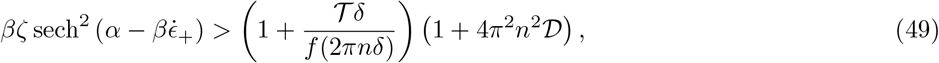

For negative *α* and positive *β* (as considered here), the left hand side increases as 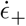 increases towards zero, and so may eventually grow sufficiently large for an ectopic instability to occur. For the parameters chosen in our numerical simulations, listed in Table I, this does not occur, but we find numerically that ectopic instabilities do occur for sufficiently large values of the tissue viscosity 𝒯 and the propagation number 𝒫. In conclusion, our model predicts that the robust formation of a single contraction in wild type embryos places constraints on the ratio of viscosities between the margin and the surrounding tissue.

**Table I:** Non-dimensional parameters used for numerical finite element simulations of the model equations Eqs. (36)-(38) (top section) and a set of corresponding dimensional quantities (bottom section) that recapitulates experimental observations.

| Parameter | Meaning | Value |
| --- | --- | --- |
| $\alpha$ | Regulation inflexion | -1 |
| $\beta$ | Regulation sensitivity | 16 |
| $\zeta$ | Regulation amplitude | 1/3 |
| $\mathcal{D}$ | Myosin diffusivity | $7.5 \times 10^{-4}$ |
| $\mathcal{P}$ | Propagation number | 2/3 |
| $\delta$ | Margin thickness | 0.208 |
| $\lambda$ | Walking kernel non-linearity | 4 |
| $\mathcal{E}$ | Margin stiffness | 1 |
| $\mathcal{T}$ | Tissue viscosity | 0.77 |
| $L$ | Margin length | 5.78 mm |
| $d$ | Margin width | 0.12 mm |
| $\sqrt{D\tau}$ | Diffusion length | 0.16 mm |
| $\tau$ | Regulation time scale | 0.5 hours |
| $\dot{\epsilon}_0$ | Maximal contraction rate | $0.33 \text{ hours}^{-1}$ |
| $\nu/E$ | Margin viscoelastic time scale | 0.5 hours |

#### F. Numerical implementation of the model

In order to compare our theoretical predictions with experiments, we implemented the model described in section II A numerically using the finite element package FEniCS 2019.1.0 [4, 5]. The numerical routine is iterative, solving the Stokes equations Eq. (31) using finite elements to obtain the instantaneous flow field ***u*** from the stress distribution ***σ***_*a*_ in alternation with explicit Euler integration of the system Eqs. (29)-(30) to update the stresses. The margin is modeled as a chain of *N* straight elements, the end points of which are advected with the flow field. *N* is chosen in order to generate margin segments with a length typical for the mesh size. For a mesh resolution of 0.1 mm, this yields *N* = 29 for an embryo half and *N* = 58 for a complete embryo. The contraction rates along each segment are calculated directly from the kinematics of the endpoints. The tension and contractility changes on the margin elements are calculated locally in a Lagrangian fashion, hence the advective terms of Eqs. (29)-(30) (which are stated in the Eulerian picture) are not included explicitly.

The mesh is auto-generated by the finite element package with a homogeneous density corresponding to at least twice the resolution of the margin segments. The mesh is created in such a way that ensures that each margin segment coincides with an edge of the mesh in order to maximise numerical accuracy. The finite elements themselves are Lagrangian third-order for the velocity and second-order for the pressure (i.e. second-order Kelvin-Hood elements), and the mesh is advected with the flow. In order to maintain a homogeneous mesh, the domain and margin are remeshed at regular intervals. Furthermore, the tension distribution *T* is interpolated linearly between the midpoints of the margin segments in order to smooth out the stress distribution ***σ***_*a*_ acting on the fluid.

In order to account for tissue growth, we also account for area changes by including a source term *γ* in the Stokes equations. These area changes are prescribed in a fashion that closely approximates the experimentally observed behavior [3], and can be decomposed into five different modes, detailed in Table II. These area changes are applied in a Lagrangian fashion, with each finite element having an appropriate value of *γ* attributed at the initialization stage which it maintains over the course of the simulation. Upon remeshing, the new compressibility values are obtained by interpolation.

**Table II:** Modes of the tissue growth term *γ*, defined in accordance with the synthetic embryo [3]. The fifth mode is time dependent with amplitude 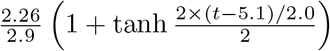. Additional parameter values taken from fit in[3]: *u*_*e*_ = 1.0488, *w*_*e*_ = 0.3956, *d*_*s*_ = 0.106, *e*_*r*_ = 1.81, *w*_*b*_ = 0.19, *r*_*s*_ = 0.8694, *w*_*s*_ = 0.1196 (all in mm), *θ*_*s*_ = 0.75 radians.

| Mode | Meaning | Value | Coefficient |
| --- | --- | --- | --- |
| 1 | EP expansion | $\frac{1}{2} \left( 1 - \tanh \left( 2(\sqrt{x^2 + (y - d_s)^2} - u_e)/w_e \right) \right)$ | 0.27/8 |
| 2 | A-P bias | $-\frac{y}{2} \left( 1 - \tanh \left( 2(\sqrt{x^2 + (y - d_s)^2} - u_e)/w_e \right) \right)$ | 0 |
| 3 | EE expansion | $1 - \frac{1}{2} \left( 1 - \tanh \left( 2(\sqrt{x^2 + (y - d_s)^2} - u_e)/w_e \right) \right)$ | 1.58/8 |
| 4 | Outer boundary correction | $-\exp \left( -0.5 \left( \sqrt{x^2 + y^2} - e_r \right)^2 / w_b^2 \right)$ | 1.06/8 |
| 5 | PS ingression | $-\exp \left( -0.5 \left( \sqrt{x^2 + (y - d_s)^2} - r_s \right)^2 / w_s^2 - 0.5 \left( \tan^{-1} (x/(y - d_s)) \right)^2 / \theta_s^2 \right)$ | see caption |

In each case the embryo is initialized with uniform tension equal to the homogeneous equilibrium value, *T* = 1 + *ζ* tanh *α*, and a sinusoidal bias in contractility *c* in the anterior, with amplitude *ζ, c* = 1 + *ζ* cos *θ* where *θ* is the polar angle measured from the posterior pole.

The boundary conditions of the finite element routine are implemented slightly differently depending on the experiments being modeled, as we explain below.

##### 1. Choice of parameter values

A list of parameters used for the simulations is given in Table I. For all of the them, we can derive quantitative, or at least quantitative constraints, from our experimental observations ([3] and this study). When constraints are qualitative, we choose parameter values such that the relevant dimensionless quantities satisfy the constrains without excessively deviating from unity. Several parameters are required to be sufficiently large to yield sufficiently strong nonlinearities in the model: the product *βζ* must be sufficiently large for spontaneous symmetry breaking, as seen in anterior epiblast halves; *λ* must be sufficiently large such that the contraction rate in the posterior is not limited by tension. The stiffness of the margin (relative to the magnitude of active tensions) is chosen in accordance with the observed elastic strains at the margin ∼ 10. Ablation experiments revealing tensions in the anterior that are comparable to tensions in the posterior suggest that *ζ* must smaller than one. Based on the rapid redirection of tissue motion in our experiments, s chosen to be under an hour. The effective myosin diffusivity *D* and the regulation time scale *τ* are constrained by the rapid redirection of tissue motion in our experiments and the estimated width of the fronts, which based on both velocity profiles and the length of supracellular cable is commensurate with the width of the margin, in the hundreds of microns. These parameters along with *α* control the extent *ρ* of the contractile domain, with values ∼ 1*/*3 (below a half) in experiments suggesting *α <* 0.

##### 2. Intact embryo

In this case the margin is fully immersed within the domain and is not subject to a boundary condition. The initial shape of the embryo is circular with radius 1.81mm, while the initial locus of of the margin is circular with radius 0.92mm and offset toward the posterior by a distance of 0.106 mm. At the outer edge of the embryo a no-slip condition is applied, and it is assumed that the embryo remains in circular shape with a radial boundary velocity *u*_*n*_(*t*). This velocity is calculated at each time step from the area changes *γ* using the divergence theorem.

##### 3. Posterior and anterior halves

In this case the boundary of the embryonic half is set by a linear cut orthogonal to the anterior-posterior (A-P) axis and through the center of the embryo proper. On the cut a no-slip condition is applied in the case of reattachment, and a no stress condition is applied in the case of no reattachment. The outer boundary moves with a velocity *u*_*n*_(*t*)***n*** that is determined as in the intact case from the total area changes. The margin has two end points which are situated on and advected with the cut, where a no flux boundary condition is applied to the contractility *c*. The no-slip condition for a reattached border is enforced through a penalty term that is large but finite, allowing a slight indentation of the border where it is pulled by the margin, as seen in experiments.

##### 4. Intact embryo with obstacle

In this case the embryo is modeled as in the intact case, except that the Stokes equation Eq. (31) is modified to include an additional friction term that is proportional to the tissue velocity and locally confined to a rectangular domain that covers the embryo fully in the lateral direction and is confined to a narrow window of twice the margin thickness, 2*d* = 0.24 mm in the A-P direction, centered on the embryo proper. The dimensionless friction coefficient is chosen sufficiently large to reduce tissue velocities by a factor of 10 in the frictional domain.

##### 5. Intact embryo with modified contractility

In this case the embryo is modeled as in the intact case, except that the initial condition is changed in order to account for transient modification of contractility through H1152 and Calyculin A. In the case of initial inhibition by H1152 in the posterior, the initial myosin profile is defined by of 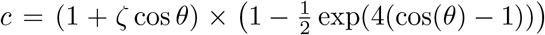. In the case of contractility enhancement in the anterior by Calyculin A, we instead modify the contractility as 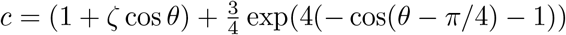.

